# Invariant neural responses for sensory categories revealed by the time-varying information for communication calls

**DOI:** 10.1101/492546

**Authors:** Julie E. Elie, Frédéric E. Theunissen

## Abstract

Although information theoretic approaches have been used extensively in the analysis of the neural code, they have yet to be used to describe how information is accumulated in time while sensory systems are categorizing dynamic sensory stimuli such as speech sounds or visual objects. Here, we present a novel method to estimate the cumulative information for stimuli or categories. We further define a time-varying categorical information index that, by comparing the information obtained for stimuli versus categories of these same stimuli, quantifies invariant neural representations. We use these methods to investigate the dynamic properties of avian cortical auditory neurons recorded in zebra finches that were listening to a large set of call stimuli sampled from the complete vocal repertoire of this species. We found that the time-varying rates carry 5 times more information than the mean firing rates even in the first 100 *ms*. We also found that cumulative information has slow time constants (100-600 *ms*) relative to the typical integration time of single neurons, reflecting the fact that the behaviorally informative features of auditory objects are time-varying sound patterns. When we correlated firing rates and information values, we found that average information correlates with average firing rate but that higher-rates found at the onset response yielded similar information values as the lower-rates found in the sustained response: the onset and sustained response of avian cortical auditory neurons provide similar levels of independent information about call identity and call type. Finally, our information measures allowed us to rigorously define categorical neurons; these categorical neurons show a high degree of invariance for vocalizations within a call-type. Surprisingly, call-type invariant neurons were found in both primary and secondary avian auditory areas.

**Author Summary:** Just as the recognition of faces requires neural representations that are invariant to scale and rotation, the recognition of behaviorally relevant auditory objects, such as spoken words, requires neural representations that are invariant to the speaker uttering the word and to his or her location. Here, we used information theory to investigate the time course of the neural representation of bird communication calls and of behaviorally relevant categories of these same calls: the call-types of the bird’s repertoire. We found that neurons in both the primary and secondary avian auditory cortex exhibit invariant responses to call renditions within a call-type, suggestive of a potential role for extracting the meaning of these communication calls. We also found that time plays an important role: first, neural responses carry significant more information when represented by temporal patterns calculated at the small time scale of 10 ms than when measured as average rates and, second, this information accumulates in a non-redundant fashion up to long integration times of 600 ms. This rich temporal neural representation is matched to the temporal richness found in the communication calls of this species.

## Introduction

Information theoretic analyses are well suited to the study of neural representation since this mathematical framework was developed to quantify and optimize the encoding of informative signals in communication channels [1]. In sensory systems, Information Theory (IT) has been applied extensively as a complimentary approach to the estimation of stimulus-response functions such as tuning curves, spatio-temporal or spectro-temporal receptive fields or other higher-level encoding models [2]. Information theoretic approaches have been particularly powerful in explorations of the nature of the neural code and its redundancy or efficiency [3–6]. For example, IT was used in early studies in the visual system to demonstrate that spike patterns contain information beyond average rate both for static images [7] and dynamic visual stimuli [8]. IT was also used to show that spike doublets can contain synergistic information that cannot be explained by an analysis of successive single spikes [9] and that, although information can only decrease in a signal processing chain, the neural coding efficiency increases as one moves to higher levels of sensory processing [10]. Finally, IT investigations also revealed that neural efficiency is higher when sensory systems process natural stimuli versus synthetic stimuli [11–13], in support of ethological theories of optimal sensory processing [14].

In sensory systems, the mutual information between a stimulus and the neural responses has often been estimated in a stimulus reconstruction framework and for continuous dynamic stimuli in stationary conditions, where time averages can be performed. In the stimulus reconstruction framework, one attempts to estimate the information about all aspects of the stimulus; for example, in audition, the stimulus would be represented by its exact sound pressure waveform. As long as the stimulus set is rich (i.e. has very large entropy), the mutual information can be an estimate of the maximum information that can be transmitted by a neural communication channel, also known as the channel capacity [3, 4]. For instance, one can obtain the mutual information of an adapted auditory neuron processing white noise or colored noise sounds [11]. As long as the stationary assumption is valid, using continuous stimuli is also beneficial as it provides large data sets that are needed to estimate the joint probability of stimuli and neural responses, both of which can have high dimensions. Even in these conditions, it is noteworthy that a direct estimation of information is only possible when many repeats of the same stimulus can be obtained [15] or when simplifying assumptions are made [9]. Ultimately, the calculation of information based on stimulus reconstruction gives a single number corresponding to the information transmitted by a single neuron or an ensemble of neurons for a particular stimulus ensemble. By repeating the calculation for different stimulus ensembles, one can investigate how the channel capacity of particular neurons or neural ensembles might depend on the stimulus statistics (e.g. for natural vs synthetic stimuli). Furthermore, by repeating the calculation using different symbols to represent the response, the potential nature of the neural code (e.g. time patterns vs. rate) can be revealed.

Here we are using IT in a different sensory encoding context: the accumulation of information in a recognition task, such as face recognition in the visual system [16] or word recognition in the auditory system [17]. Recognition or identification is one of the key computations performed by higher sensory areas as opposed to the task of efficient stimulus representations that is performed in lower sensory areas and that might therefore be well quantified by information values based on stimulus reconstruction. In the recognition tasks, each stimulus is described by a simple label, such as the word corresponding to a given speech sound or that is used to label a given visual object. The relevant value of information in that task is then the information about these discrete labels and the information capacity of the system in its ability to identify the stimulus as a whole. In the recognition framework, one can ask how the information about the stimulus identity or label changes as a function of time relative to the stimulus onset and to what extent that time-varying information is redundant and, thus, how it accumulates over time. For example, one could ask at what time after stimulus onset does the performance of single neurons or ensemble of neurons match a behavioral performance of word recognition. Such an IT analysis has been performed in the primate visual system using a delay-matching to sample paradigm, and using spike counts, estimated in progressively longer windows, as the neural symbols [18].

In information studies based on continuous stimulus reconstruction, the neural code can be investigated in terms of its temporal resolution (i.e. letter size) and its integration time (i.e. word length). While the same properties of the neural code can be deciphered in the recognition framework, one can also examine the relationship between spikes at different points in time and time-varying information. This analysis is meaningful because a time zero corresponding to stimulus onset can be clearly defined and is behaviorally relevant. Moreover, stimulus-response functions for such discrete stimuli are not time-invariant. Responses in sensory neurons, in vision [19, 20] and in audition [21, 22], are often characterized by an onset response (or on-response) and a sustained response, where both the precision of spikes and the information coded might be different [23, 24]. For example, a first spike latency code has been proposed as a fast encoding scheme in vision [25], audition [26] and somato-sensation [27]. Rolls et al. tested this hypothesis, by quantifying the fraction of information that is present in the first spike relative the on-going response [28].

Finally, in the recognition framework, one can also compare the information values obtained when different labelling schemes are used for identifying the stimuli as objects. For example, speech sounds could be labeled hierarchically as unique utterances, as phonemes, as syllables, as words, etc.. One can then compare time-varying information about each of the levels in such hierarchical labelling scheme and gain insight on the neuro processing involved in object categorization. Although such a hierarchical representation of stimulus features has been used in encoding models for studying human processing [29], it has not yet been used in an IT analysis.

In this study, we developed a new approach for estimating time-varying information and cumulative information for sensory object identification task. Our approach assumes that time-varying neural responses can be modeled as inhomogeneous Poisson processes and generalizes well to large number of stimulus categories and to long integration times relative to the dynamics of the time-varying response. Our motivation for developing this methodology was to gain additional understanding on the neural representation of communication signals in high level auditory areas. Animal communication calls, just as speech sounds in humans, are categorized into behaviorally meaningful units. Significant progress has been made in identifying brain regions involved in categorizing sounds, in particular in the primate brain, where neural responses that are correlated with progressively more abstracts concepts are found in primary auditory cortex, the lateral belt of the auditory cortex and the prefrontal cortex [30]. However, the neural computations involved in generating categorical responses remain poorly described [31] and only a small number of studies have examined the neural categorization of natural communication calls in non-human species [32–36]. We and others have been developing an avian model system to study the neural processing of relatively large and complex vocal repertoires [37, 38]. Our prior studies include a detailed bioacoustical analysis of the features that define each call-type of the complete vocal repertoire of the zebra finch [39] and the first characterization of neural responses to the calls from that large repertoire in primary and secondary avian auditory cortical areas [40]. In that study, we found that approximately 45% of auditory neurons encode information about call-type categories. Among those, a minority show strong selectivity for single call-type categories and invariance for calls within that category. Here, we investigated the processing in time that could lead to those observed categorical responses by comparing the time-varying information for stimuli labelled as individual utterances to the time-varying information for the same stimuli labelled by their call-type category. With that analysis, we were able to obtain values of temporal integration for stimulus identification and call-type category identification. We also analyzed the relationship between the time-varying firing rate and the time-varying information and, in particular, examined differences in selectivity in the onset versus sustained response. Finally, we used anatomical data to examine the distribution of neurons in primary and secondary avian auditory cortical areas with distinct responses properties as revealed by this IT analysis.

## Results

We studied the time-varying information in a population of neurons recorded from primary and secondary regions in the avian auditory cortex of head-fixed urethane anesthetized zebra finches listening to a large set of natural communication calls. Zebra finches emit various calls in different behavioral contexts and have a complete vocal repertoire composed of 11 call-types. The acoustical characteristics of each call-type have previously been described in detail [39]. Here we focused on the neural representation of 9 call-types: 1) 3 pro-social calls emitted for pair bonding and social cohesion: the Distance Call (DC), the Tet Call (Te), the Nest Call (Ne); 2) the Song (So) that is emitted as a sexual display in males; 3) 2 calls emitted in aggressive encounters: the aggressive Wsst Call (Ws) and the Distress Call (Di); 4) 1 alarm call, the Thuk Call (Th) and 5) 2 calls emitted by juveniles: the Begging Call (Be), used by young to request food, and the Long Tonal Call (LT), a contact call that is a precursor of the adult DC. A stimulus set was composed of approximately 10 different exemplars of calls, train of calls or song produced by different vocalizers for each of the 9 call-types. The call stimuli were randomly sampled from a large annotated data base of calls and songs from the complete repertoire of the male and female zebra finch. Neural responses were recorded using electrode arrays implanted in both hemispheres of 4 male and 2 female adult zebra finches. We recorded from a total of 914 single auditory units in both primary (Field L) and secondary (CML, CMM and NCM) avian auditory areas. In previous analysis, we showed using a decoding approach that information about call-types was found in 404 (44%) of these units [40]. Here, we further restricted our population analysis of information to neurons, from that same data set, that reached a significant level of information about the stimulus (see methods) and analyzed the time-varying information of 337 neurons during the first 600 ms of their response. Note that many calls are shorter than 600 ms, but also that they are often produced in succession with short inter-call intervals. Thus, this analysis window could contain two or more calls or the beginning of a longer song motif comprised of multiple syllables. We only analyzed the response in the first 600 ms because the estimation of the cumulative information for longer time windows became unreliable, as we will explain below. Additional details on the neurophysiological recordings can be found in the methods section and in Elie and Theunissen [40].

In the *Results*, we first describe the approach we developed to estimate instantaneous and cumulative time varying information. We illustrate these calculations with specific examples of model and actual neurons in the avian auditory cortex. We then analyze the time-varying coding properties of the avian cortical auditory neurons with an emphasis on: the relationship between spike rate and instantaneous information, the time constants observed for the cumulative information and the relative fraction of stimulus cumulative information that is used for extracting the behaviorally relevant categories corresponding to distinct call-types.

### Estimation of the time-varying information

At a given time t, the *instantaneous* mutual information between the stimulus S and the response *Y_t_* can be written as a difference in Shannon entropies:

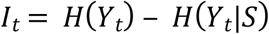

Here, *H*(*Y_t_*) is the response entropy for a window at time *t*, while *H*(*Y_t_|S*) corresponds to entropy of the response given the stimulus or the conditional response entropy. *H*(*Y_t_*|*S*) can also be called the neural noise since it represents the variability in the neural response to the same stimulus. For spiking neurons, *y_t_* represents the number of spikes in the window at time *t* (Note: In our notation, capitals are used for random variables and lower case for a sample from that random variable).

Similarly, the cumulative mutual information in neural responses that are discretized into time intervals is given by:

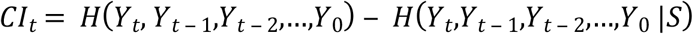

The entropies now include the time course of the neural responses starting at *t*=0 and up to time *t*. The reference time *t*=0 is set to the stimulus onset in our analyses but could be any arbitrary reference point.

The conditional response entropy and the response entropy are obtained from the distribution of the conditional probability of neural responses given the stimulus, *p*(*y_t_*|*s*), and the distribution of probability of each stimulus *p*(*s_i_*):

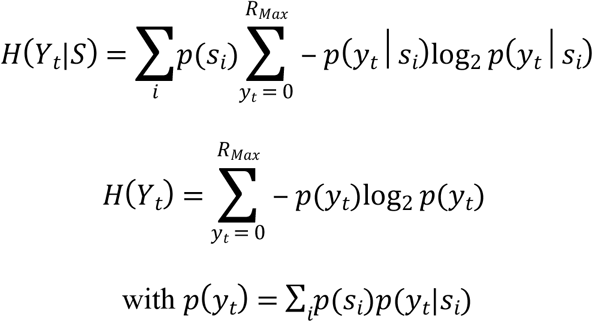

*y_t_*, the neural response at time *t*, is measured in spike counts and takes values from zero to a maximum rate value, *R_Max_*(for example, as dictated by the neuron’s refractory period or numerically as *p*log *p* becomes infinitely small for high spike counts that have very small probability of occurring). The probability of the stimulus *p*(*s_i_*) is usually taken as 1/*n_s_*, where *n_s_* is the number of stimuli, unless the study incorporates natural stimulus statistics. The probability of spike counts at time *t* given a stimulus, *p*(*y_t_*|*S_i_*), could be estimated empirically by recording hundreds or thousands of responses of the same neuron to the same stimulus. Although this approach has been shown to be possible in certain preparations [15], it severely limits the number of stimuli that can be investigated in most neurophysiological experiments. Here we propose a parametric approach where we model the distribution of neural responses to a given stimulus *s_i_* as an inhomogeneous Poisson process. The conditional probability of response (spike count) given the stimulus is then:

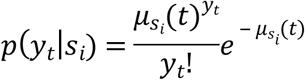

where *μ_S_i__*(*t*) is the mean response at time *t* for stimulus *s_i_*. This mean rate was estimated empirically using the time varying kernel density estimation (KDE) proposed by Shimazaki and Shinomoto [41]. The instantaneous information estimated in this fashion is relatively straightforward, as long as the Poisson assumption is valid and a sufficient number of trials is obtained to estimate *μ_S_i__*(*t*) (see methods).

The estimation of the cumulative information then extends this approach to joint probabilities of responses in successive time windows, (*y_t_, y*_*t*−1_,*y*_*t*−2_,…). Due to what has been labelled as the “curse of dimensionality”, the numerical estimation of the unconditional probability becomes exponentially more expensive as the integration time increases. We evaluated multiple approaches based on different assumptions and found that Monte Carlo with importance sampling yielded the best results (see methods and Sup. Fig. 3 for comparison to alternative approaches).

Given our Poisson assumption, the conditional probability of response at *t* is independent of the conditional response at previous times.

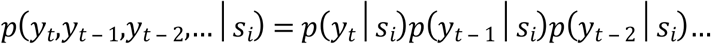

Because of this probabilistic independence, it can be shown that the joint conditional response entropy is simply the sum of the conditional response entropies at each time point (see methods):

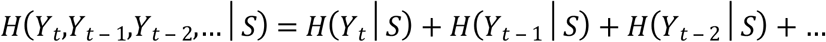

Thus, the estimation of the conditional response entropy is straightforward and not affected by the integration time. The problem of dimensionality arises in the estimation of the unconditional probabilities of response and the corresponding response entropy.

The probability of the time varying response is the joint probability of observing (*y_t_, y*_*t*−1,_*y*_*t*−2_,…). This joint probability cannot be expressed as the product of the probabilities at different times because these are not independent. More intuitively observing a particular *y*_*t*−1_ will affect the probability of observing *y_t_*. This is true because the time varying means of the Poisson distributions *μ_S_i__*(*t*) are correlated in time; for example, if *μ_s_i__*. (*t* − 1) is high we might expect *μ_S_i__*(*t*) to also have high values. These high values could be true for one particular stimulus *S_i_* but not for the other stimuli. Then observing a high value of *y*_*t*−1_ would predict a higher value than expected for *y_t_* (and an increase in probability that it was caused by *S_i_*). The joint unconditional probability distribution is:

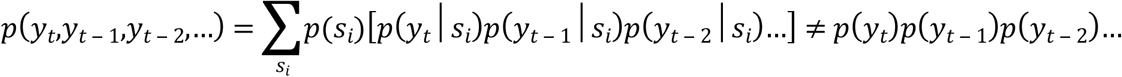

Given the lack of independence, the response entropy must then be calculated from the joint probability distribution:

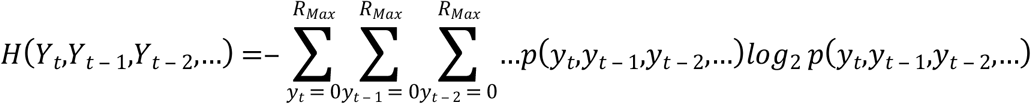

The estimation of this entropy was performed using Monte Carlo with importance sampling. In Monte Carlo, random samples of a vector (*y_t_, y*_*t*−1,_*y*_*t*−2_,…) are drawn from a proposal distribution q(*y_t_, y*_*t*−1_,*y*_*t*−2_,…) and used to estimate the expected value of log_2_ *p*(*y_t_,Y*_*t*−1_,*y*_*t*−2_,…) by an algebraic average weighted by the likelihood ratio of *p*(*y_t_, y*_*t*−1_,*y*_*t*−2_,…)/*q*(*y_t_, y*_*t*−1_,*y*_*t*−2_,…). The sampling stops when entropy estimations reach an equilibrium. Information estimations are also known to suffer from positive bias [42]. Here, biased-corrected estimates and errors on information values were obtained from Jackknifing the estimation of the time-varying rates and bootstrapping the Monte Carlo samples.

For these calculations, one has also to determine the size of the time window used for estimating the instantaneous information and correspondingly the steps for the cumulative information. This time window is used to estimate spike counts and the average rate *μ_S_i__*(*t*) and depends both the dynamics of the stimulus and on the response properties of the neurons. By performing a coherence analysis on spike trains and a power spectral analysis on time varying rate in response to natural stimuli, we found that 10 ms (or 50 Hz) captured between 97% and 99% of the dynamics in our system (see methods and Sup. Fig. 1).

Finally, we also estimated the information values for stimulus categories by performing the weighted sum of probabilities for stimuli belonging to each category. The information about categories at time *t*:

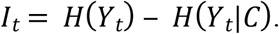

is obtained from the conditional probability of response given the category *c_k_*, which is in turn calculated as the average conditional probability of response for the stimuli belonging to that category:

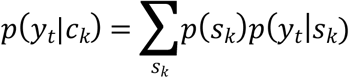

Here, *p*(*s_k_*) is the probability of occurrence of stimulus *s_k_* within the category *c_k_*. In controlled playback experiments (as here), *p*(*s_k_*) is 1/*n_S_k__* where *n_S_k__* is the number of stimuli used to sample category *c_k_*. Similarly, *p*(*c_k_*), the probability of occurrence of vocalizations in category *c_k_* was taken as 1/*n_c_* where *n_c_* is the number of categories. In our system, the stimuli are individual renditions of vocal communication calls that fall into 9 call categories of the zebra finch vocal repertoire. We will contrast stimulus information to categorical information, both instantaneous and cumulative for these behaviorally relevant categories of call-types.

### Time-varying information for model neurons

To validate our approach and to illustrate the behavior of time-varying information values, we calculated instantaneous and cumulative information for model neurons with simple and stereotyped response properties. Figure 1 shows the firing rates, raster plots and information values for 3 model neurons in response to 4 stimuli (S1-S4). One model neuron responds to the 4 stimuli with different mean firing rates that are constant in time (Rate Neuron). A second model neuron responds to the four stimuli with the same fixed firing rate but with different latencies (Onset Neuron). The third model neuron also responds with equal average firing rates to the four stimuli but the response occurs at different times (Temporal Neuron).

**Figure 1.**
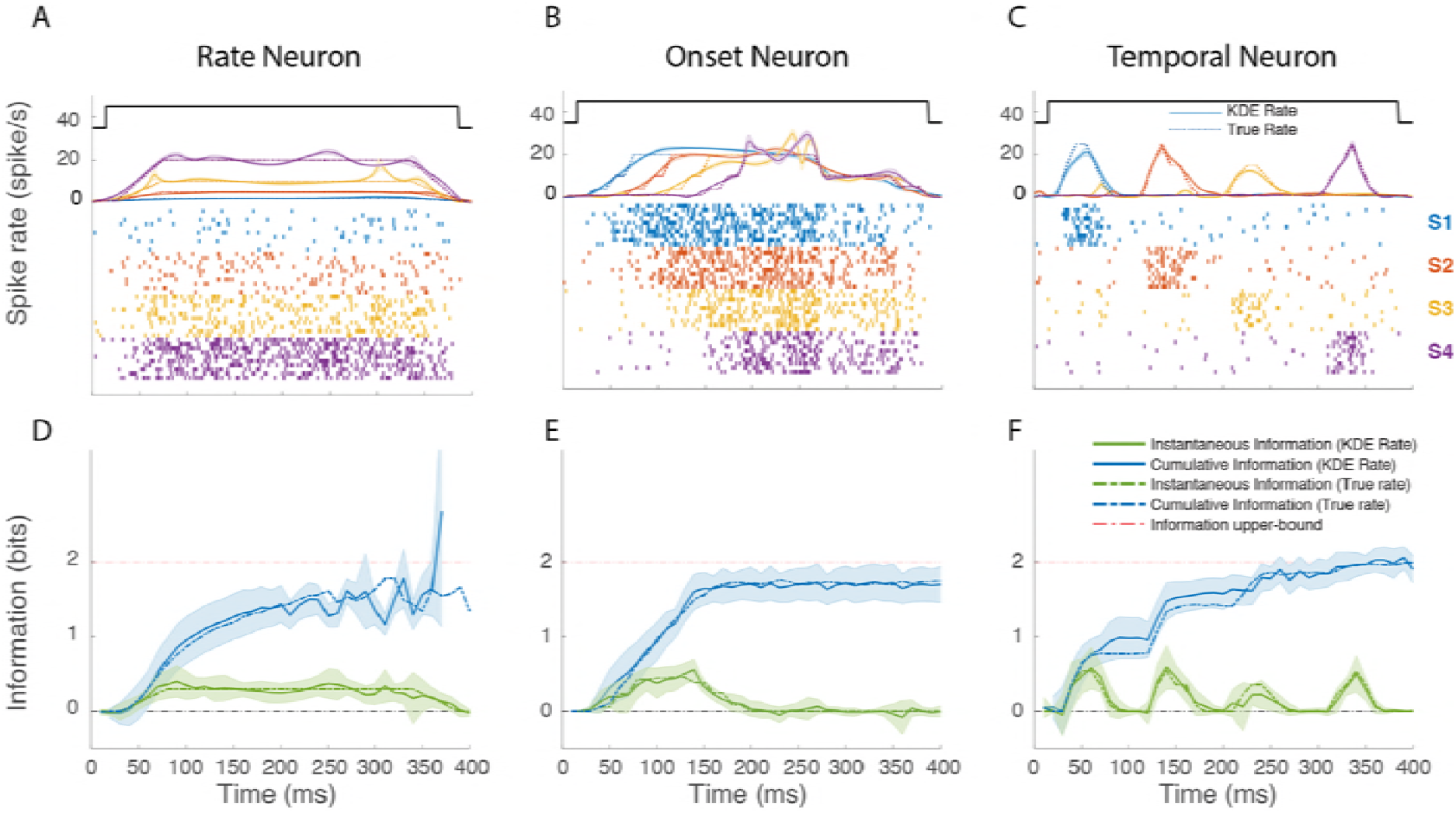
Instantaneous and Cumulative Information for 3 Model Poisson Neurons. The *top row* (**A,B,C**) shows the firing rate and simulated spike rasters of 3 model neurons to 4 hypothetical stimuli (S1-S4). The onset and offset of all stimuli are identical and shown by the top solid black line. The model rates for the four stimuli are shown in colored dotted lines. These rates are used to generate spike patterns using an inhomogeneous Poisson process. Ten realizations for S1-S4 are shown as spike rasters. The rate recovered from those rasters using a time-varying kernel density estimate (KDE) are shown as colored solid lines. The shaded area around each line indicate the error on the KDE rate estimated by the standard error on the Jackknife estimates. The *bottom row* (**D,E,F**) shows the instantaneous and cumulative information obtained using either the true model rate or the recovered rate from KDE. The error bars in the instantaneous information calculations were obtained by jackknifing the KDE estimates. For the cumulative information calculations, the Monte Carlo simulations were repeated for each jackknife estimate. The error bars therefore include both the errors in the estimation of the true rate from spike rasters (data limitation) and the errors in the estimation of the cumulative information due to limited sampling (computational limitation). The time-varying response rates were chosen to illustrate neural representation for stimuli using different neural codes labelled here as *Rate* (**A, D**), *Onset* (**B, E**), and *Temporal* (**C, F**).

These simulations illustrate some very basic principles of neural coding. First, many different response profiles can lead to very similar rates of information: in all three simulations, the cumulative information approaches the maximum possible value (2 bits). Second, the coding capacities of neurons are a function of both the range of firing rates that can be achieved (as in the rate neuron) and the modulations in time of this neural activity. Third, the estimation of the instantaneous information gives an incomplete picture of the neural coding of a neuron as it does not incorporate the redundancy or independence of the neural representation over time. For example, on the one hand, comparing the instantaneous information in the Rate neuron to that of the Onset neuron, one might erroneously conclude that the Rate neuron has more information while, in fact, the cumulative information shows that the Onset neuron is more informative at short time scales. On the other hand, one can also observe that the cumulative information in the Rate neuron continues to increase while the firing rate is constant; additional time points allows for a better assessment of that firing rate by time-averaging out neural noise.

These simulations also allowed us to validate our methods. The KDE for the empirical estimation of the time-varying rate based on the generated spike rasters (solid line) gave very good predictions of the actual model rates (dashed lines): over all stimuli and model neurons, the average error was less than 0.02 spikes/ms. Not surprisingly then, using the actual rate versus the estimated rate yielded practically identical results in the information calculations (dashed vs solid lines). We also checked that the bias corrected estimates were accurate: the instantaneous information was indeed centered at zero when the response to the 4 stimuli was identical. We verified that the actual values of instantaneous and cumulative information were correct. For example, in the Temporal neuron a peak instantaneous information of 0.5 bits is expected as 1 out 4 stimuli will be almost perfectly discriminated. Finally, we also assessed the limitations of Monte Carlo with importance sampling for the estimation of the cumulative information. For neurons, with continuously high firing rates, this estimation can become unreliable at longer integration times as illustrated by the calculation for the Rate neuron. However, in those cases, the estimate of the standard error also increased drastically and allowed us to define end points for the calculation of the cumulative information.

### Avian auditory neurons have Poisson statistics

Since our estimation of the time-varying information values was based on a Poisson parametrization for the distribution of spike counts, we first assessed the validity of this assumption. Given the small number of trials, we could only assess whether the first and second moments (mean and variance) obeyed Poisson statistics. Figure 2 shows the Fano Factor estimated at consecutive time points after stimulus onset and both for individual stimuli (onset) and average across stimuli. The mean Fano factor for the population was 1.04 with SEM of 0.04.

**Figure 2.**
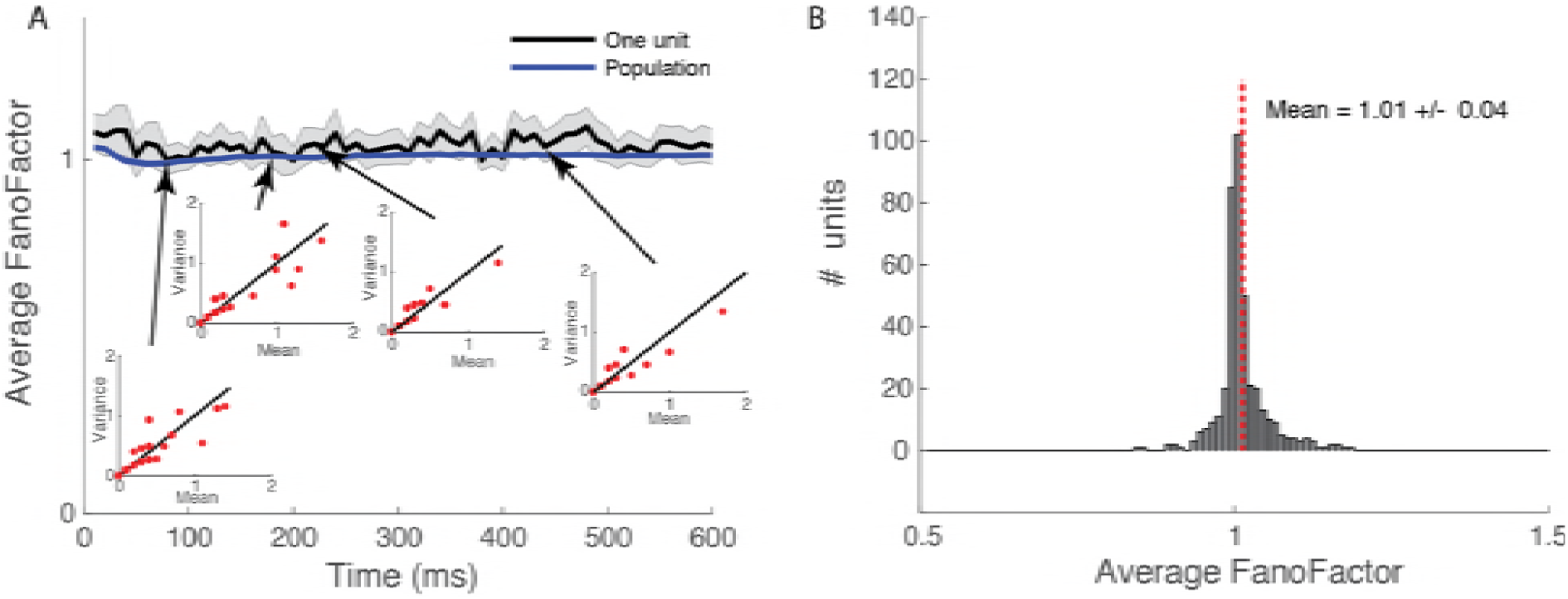
Avian Auditory Neurons have Poisson Statistics. **A**. Time-varying Fano Factor (Variance/Mean) obtained from the empirical distribution of spike counts estimated in successive 10 ms windows. Spike count distributions are obtained from 10 trials and the average Fano Factor from repeating the estimation of mean and variance calculation for all stimuli presented (min=54, max=104). The Fano Factor is shown for one example neuron (solid black line) and also averaged across the entire population (n=404). Error bars are ± two SEMs. The insets show the mean and variance spike count for the example neuron at a given time shown by the arrows. On those plots, each red dot corresponds to one stimulus (many points overlap). **B**. Distribution of time averaged Fano Factors for the population of neurons (n=404). Fano Factors are not significantly different from 1.

### Time-varying Information for 3 example neurons

In Figures 3-5, we show the time-varying firing rates, the time-varying instantaneous information and the cumulative information for 3 zebra finch auditory neurons with distinct response properties. The neuron in Figure 3 responded robustly to all communication calls with high and reliable firing rates. It also responded in a time locked fashion sometimes at multiple time points for single calls. Although this neuron was not selective for a particular stimulus or call-type category, by combining rate and temporal codes it reached very high levels of instantaneous information. Moreover, this instantaneous information showed little redundancy yielding very high cumulative information. One can also observe, that for this particular neuron, the average time-varying firing rate is not correlated with instantaneous information. This neuron shows a strong onset response in its firing rate while the instantaneous information is almost constant and even slightly lower during the onset response. The neurons in Figures 4 and 5 have much lower firing rates and exhibit selectivity for a call-type, the Distance Call (DC) and the Wsst Call (Ws) respectively. The neuron in Figure 4 exhibits both an onset and sustained response both of which are selective for DC. The neuron in Figure 5 has a much longer latency for response with correlated peaks in firing rate and instantaneous information found between 100 and 300 ms after stimulus onset (Example of spike rasters from single trials for these three neurons are shown in Sup. Figs. 4–6).

**Figure 3.**
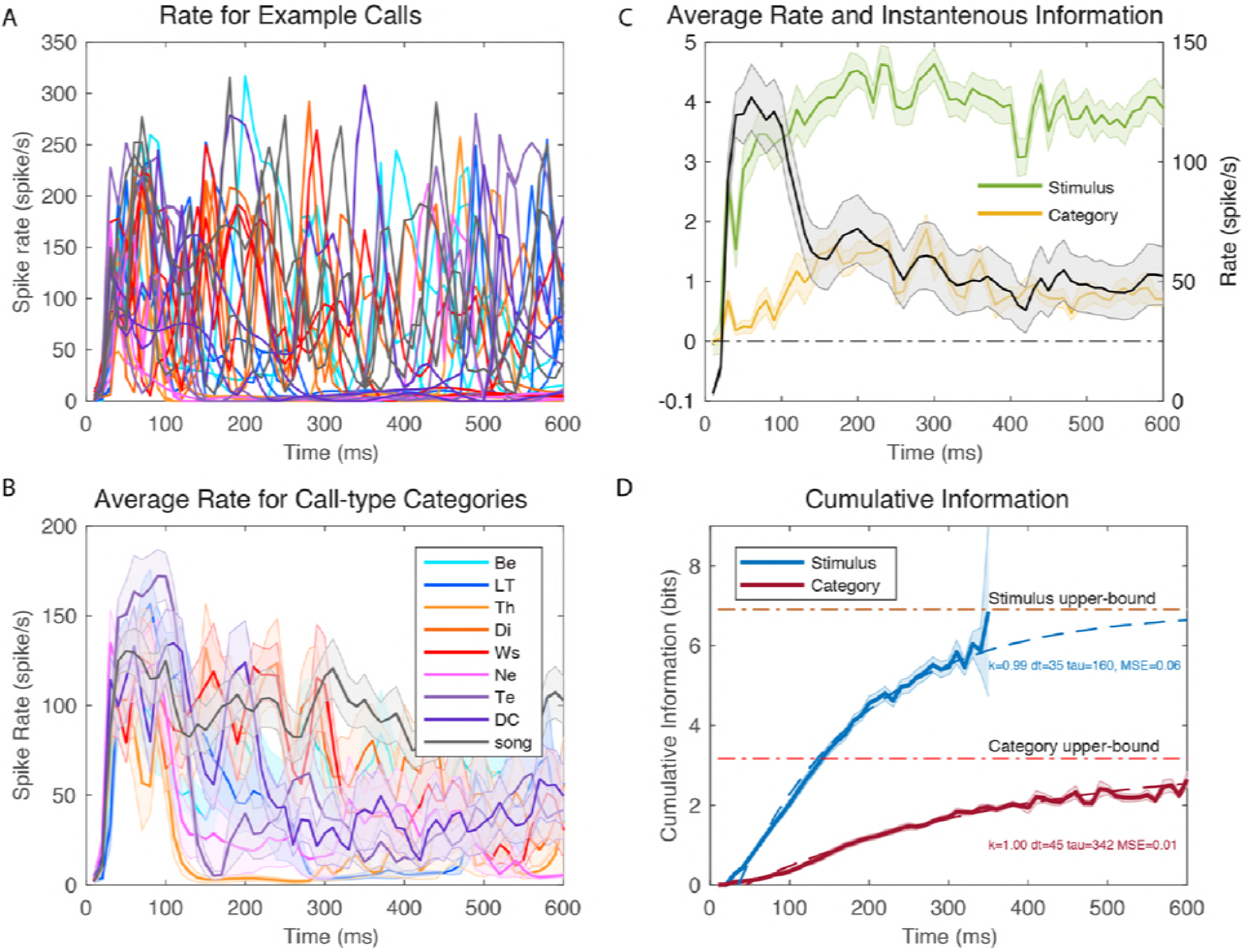
Time-varying Firing Rates and Information Values for Example Neuron 1. **A**. Time-varying firing rate obtained from kernel density estimation (KDE) of ten repetitions of different example stimuli. The plot shows the responses to two examples of calls from each call-type: 18 time series color coded by call-type using the color scheme shown in the legend of B. 0 ms corresponds to stimulus onset. **B**. Averaged time-varying firing rate obtained for each call-type category. These average responses are obtained by averaging the KDE estimates of firing rates for each stimulus belonging to the same call-type (~ 10 example stimuli per call-type). Shaded error bars show ± one SEM. **C**. The overall (averaged over all stimuli) time-varying firing rate for the same example neuron is shown by the solid black line (right y-axis) with ± SEM as shaded error bars. The green line and yellow line (left y-axis) correspond to the instantaneous information for stimuli and call-type categories calculated in successive 10 ms windows. Shaded error bars were obtained by a jackknife procedure **D**. Cumulative information for stimuli and call-type categories for the same example neuron. The shaded error bars are ± SE obtained by jackknifing and resampling (see Fig. 1 and Methods). Each cumulative information curve is also fitted using an exponential function (dashed lines) characterized by three parameters: a latency (dt, in ms), an exponential rate (tau, in ms) and the infinite time limit value expressed as a fraction of the maximum information achievable (k). The MSE is the mean square error of the fit in bits. This example neuron had a very high stimulus evoked mean firing rate with rapid and reliable time-varying dynamics leading to high information rates. Spontaneous firing was very low (~0 spikes/s). Spike rasters for this neuron are shown on Sup. Fig. 4.

**Figure 4.**
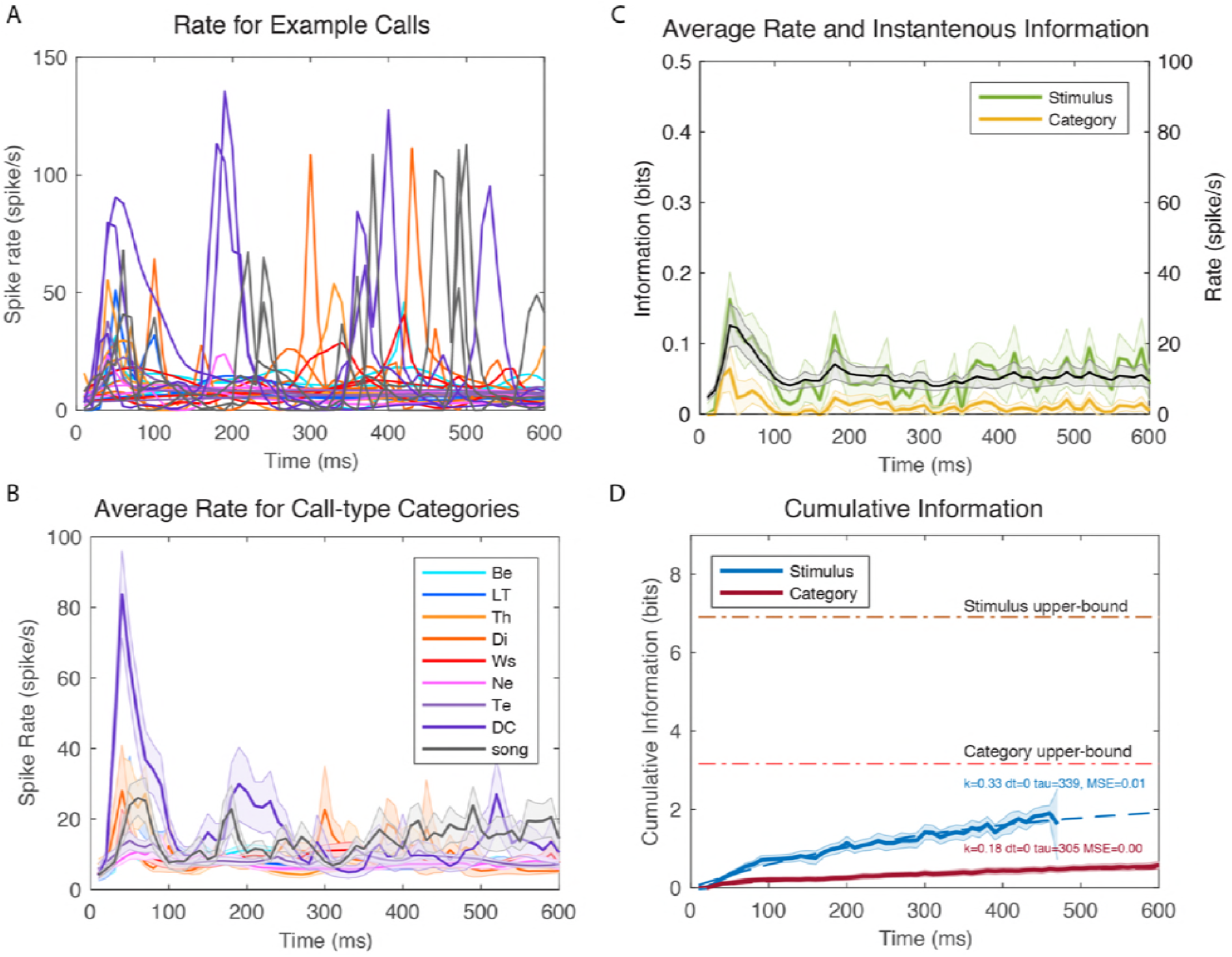
Time-varying Firing Rates and Information Values for Example Neuron 2. As in figure 3 but for a neuron with lower firing rates and displaying selectivity for Distance Calls. Spike rasters for this neuron are shown on Sup. Fig. 5.

**Figure 5.**
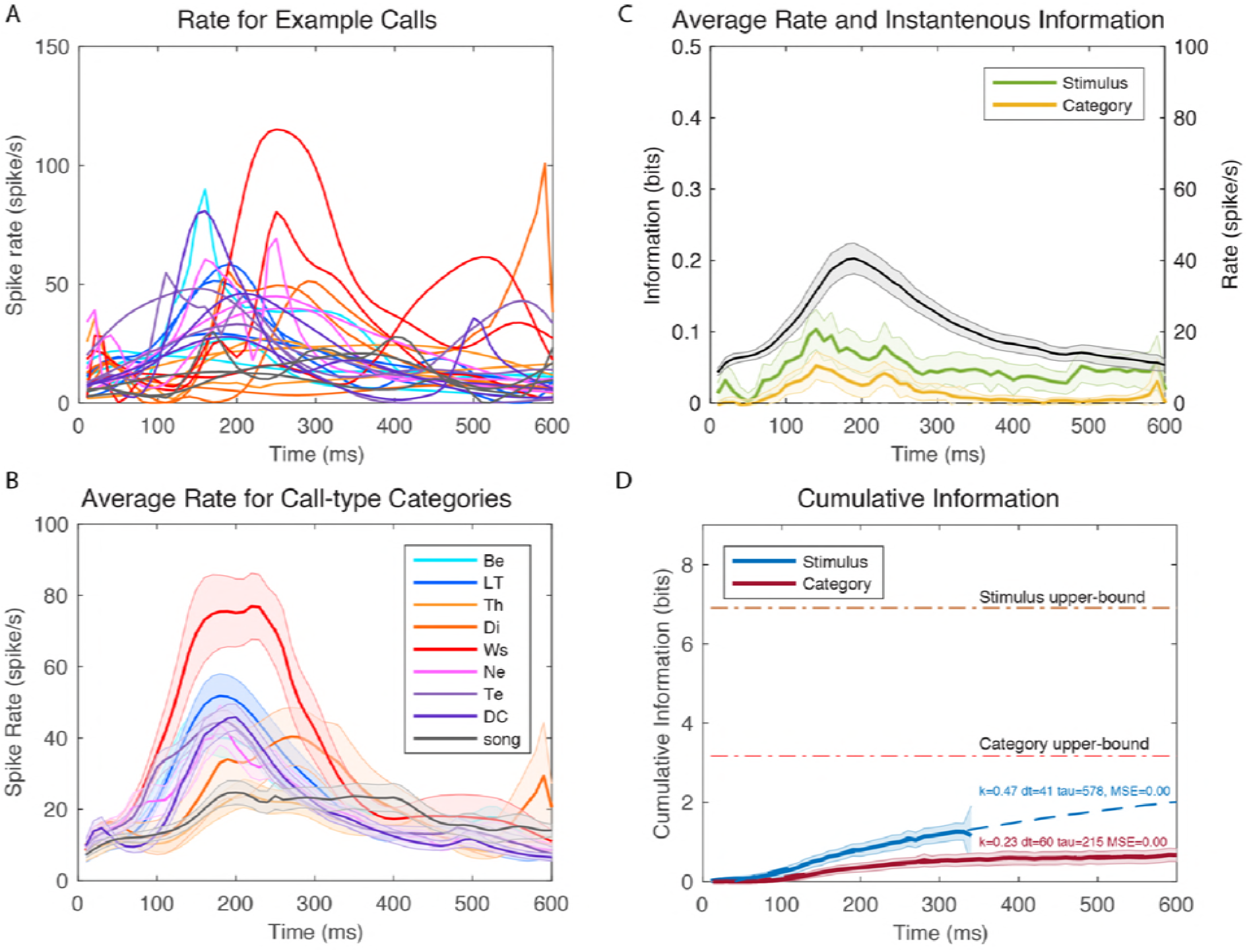
Time-varying Firing Rates and Information Values for Example Neuron 3. As in figure 3 but for a neuron with intermediate firing rates and displaying selectivity for Wsst (Ws) or Aggressive Calls. Note also the longer latency relative to the neuron shown in 3 and 4. Spike rasters for this neuron are shown on Sup. Fig. 6.

The cumulative information curves were fitted with an exponential function that allows us to quantify the time constant of information accumulation (*τ*), the saturation level (*k*) relative to the maximum information that could be achieved (*I_Max_*) and the latency (*dt*):

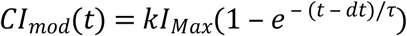

Most of the calls in the zebra finch repertoire have durations that are shorter than 300 ms [39]. In order to investigate the cumulative information accumulated at this behaviorally relevant fixed point in time, we estimated the relative cumulative information as the value of the model at 300 ms relative to I_Max_.;

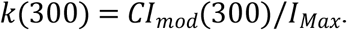

The results of the fit are shown in dashed lines in the cumulative information plot for each neuron. One can observe that the neuron in Figure 3 has exceptionally high values of saturation: with sufficiently long integrations time, single spike trials could be used to perfectly assess what stimulus (out of those used in the experiment) was hear and, thus, which call-type category it belonged to. The selective neurons in Figures 4 and 5 have much lower saturation levels as expected since they mostly respond to stimuli from a single call-type category. The neuron in Figure 5 has longer latency in its categorical cumulative information but this is not the case for the neuron in Figure 4 that has a rapid yet selective onset response. The high firing neuron (Figure 3) also has a faster time constant *τ* for the stimulus information than the selective neurons shown in Figures 4–5. All three neurons have similar and relatively slow time constants for the categorical information in the 200-300 ms range.

### Rate and Time-Varying Information: Population Analysis

Figure 6A shows the time course of the firing rate averaged across all stimuli and its relationship with the instantaneous information. On average, avian cortical auditory neurons show an onset response followed by a sustained response as observed at many stages of auditory processing and in many sensory systems. The instantaneous information, however, remains almost constant during the entire time. This is true both for the information about individual stimuli or the information about categories. Thus, the onset response is only less selective/informative than the sustained response in bits/spike but not in bits/s; when processing natural vocalizations, the onset and sustained response are equally informative. Moreover, the information in the sustained response continues to provide new information as reflected by the continuous and relatively fast increase in cumulative information shown in Figure 6B. That rapid rate of increase should be compared to the one observed when the mean firing rate for each stimulus estimated across the entire time-window was used in the calculation (dashed lines in Fig. 6B). As seen in those curves, the increases in the cumulative information for the rate code is much smaller (the information still increases because of noise averaging as explained above). The time-varying rates observed in neural responses (as illustrated in Fig. 3A) provide additional information. How much more? For the coding of individual stimuli (comparing the dashed blue line to the solid blue line in Fig. 6B), a fixed rate code (or assumption) captures only 24% of the information at 100 ms and 21% at 300 ms. The effect for categorical information is smaller because some of the coding dynamics in time-varying responses to stimuli belonging to the same category effectively become neural noise: for categorical information, a fixed rate code captures 50% of the information at 100 ms and 41 % at 300 ms.

**Figure 6.**
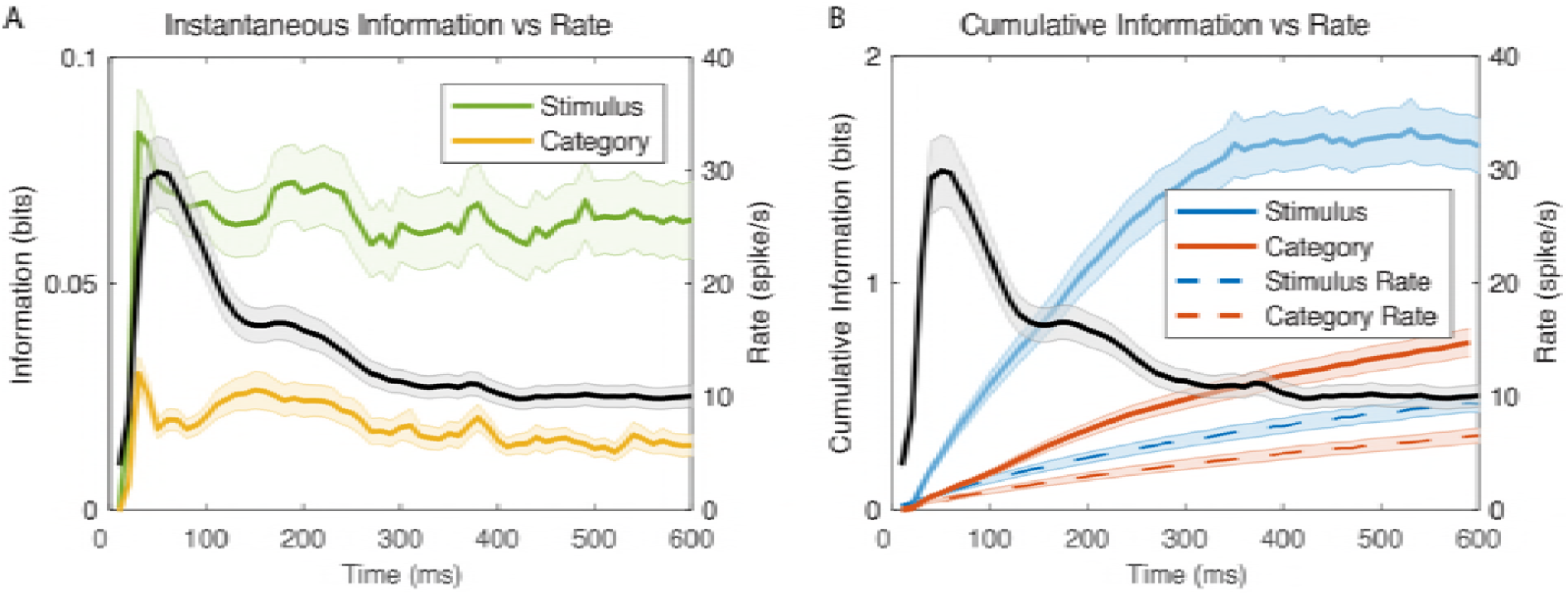
Time-varying Firing Rates and Information Values for the Population. **A**. The solid black line (right y-axis) shows the time-varying firing rate averaged across all neurons (n=337) and all stimuli played. The shaded error bars are ± SEM. The green and yellow lines (left y-axis) show the instantaneous information for stimuli and call-type categories respectively. The shaded error bars are ± SEM. **B**. The time-varying mean firing rate is shown as in A. The blue and red solid lines show the population average cumulative information for stimuli and call-type categories respectively. The blue and red dashed lines show the corresponding average cumulative information values calculated with constant averaged firing rates for each stimulus for each neuron over the first 600ms after stimulus onset (a rate code). The shaded error bars are ± SEM.

The distributions of time constants, *τ* for the cumulative information for stimuli and call-type categories are shown on Figure 9. The distributions of relative cumulative information at 300ms (k(300)) are shown on Figure 10.. The range of time constants observed across the population of neurons was large (100 ms-600 ms) with average time constants of 459 ms for cumulative stimulus information and 372 ms for cumulative categorical information. This difference in means of time constants is statistically significant (Paired t-test t(214)=3.49, p=0.00058); the ongoing time-varying rate changes continue to provide more information for decoding stimuli and less so for decoding categories of stimuli. There is also a wide distribution of relative cumulative information values (k(300) ranging from close to zero to 0.6. On average across neurons, the k(300) was 0.2 (or 20%) for stimuli and 0.15 (or 15%) for categories. These differences in relative information values are highly significant suggesting that, single neurons, capture more variability in stimuli than in categories. Note however that measures of relative information depend on the number of stimuli or categories. Therefore, a direct statistical comparison is not warranted. The comparison between stimulus representation and category representation requires estimations of expected values of categorical information given stimulus information which is performed below. Relative cumulative information values can, however, be used for stimulus and categories independently to assess other coding properties: although average time-varying rates and instantaneous time varying information are not well correlated within single neurons (as shown in Figs. 3 and 6), the relative cumulative information is correlated with average firing rates. For cumulative stimulus information, one finds an increase in k(300) of 1% per spike/s (Adj R^2^=0.36, F(1,213)=120, p=1.82 10^−22^) and, for cumulative categorical information, an increase in *k*_300_ of 0.8% per spike/s (Adj R^2^=0.34, F(1,213) = 112, p = 2.26 10^−21^).

### Analysis of the Categorical Information

We are interested in identifying neurons that could play an important role in categorizing vocalizations. We had previously identified example neurons that were highly selective for particular call-types and showed a high degree of invariance such as those shown in Figures 4 and 5 [40]. Here, we attempted to quantify the neural invariance for call renditions within call-type categories along time. For this purpose, we computed a Categorical Information Index (CII). The CII compares the actual categorical cumulative information to three potential values: a floor or minimum value (set at 0), an expected value for shared information between stimuli and categories (set at 1) and a ceiling or maximum value (set at 2). The floor is the categorical information that one would obtain if stimuli are randomly grouped. The shared-information value is the information that one would obtain if the information about stimuli is equally shared across all stimuli and the neural responses for stimuli are perfectly sorted for each natural call-type category; for example, at a given point in time the 10 renditions of DC would elicit the 10 highest rates, the 10 renditions of LT call the next 10 higher firing rates and so forth. This shared-information value does therefore assume that, for a given neuronal signal to noise ratio, neural responses segregate categories maximally while also preserving the maximum discrimination between stimuli within categories. The ceiling value is the categorical information value that one would obtain if a maximum amount of information about stimuli was used for the discrimination of categories and a minimum for discriminating stimuli within categories; it assumes maximum invariance to variations within a category. The ceiling value is equal to the stimulus information until it reaches log_2_ (*n_c_*), where *n_c_* is the number of categories, corresponding here to the 9 call-types.

Figure 7B shows the time-varying floor (dashed-green), shared (dashed-orange) and ceiling (dashed-red) values of cumulative categorical information along with the actual stimulus (solid blue) and categorical (solid red) cumulative information for 3 neurons chosen to illustrate CIIs that are below 1, around 1 and above 1. Figure 7A shows cartoon probability distributions of neural responses for particular stimuli and particular categories that correspond to the floor, shared and ceiling values.

**Figure 7.**
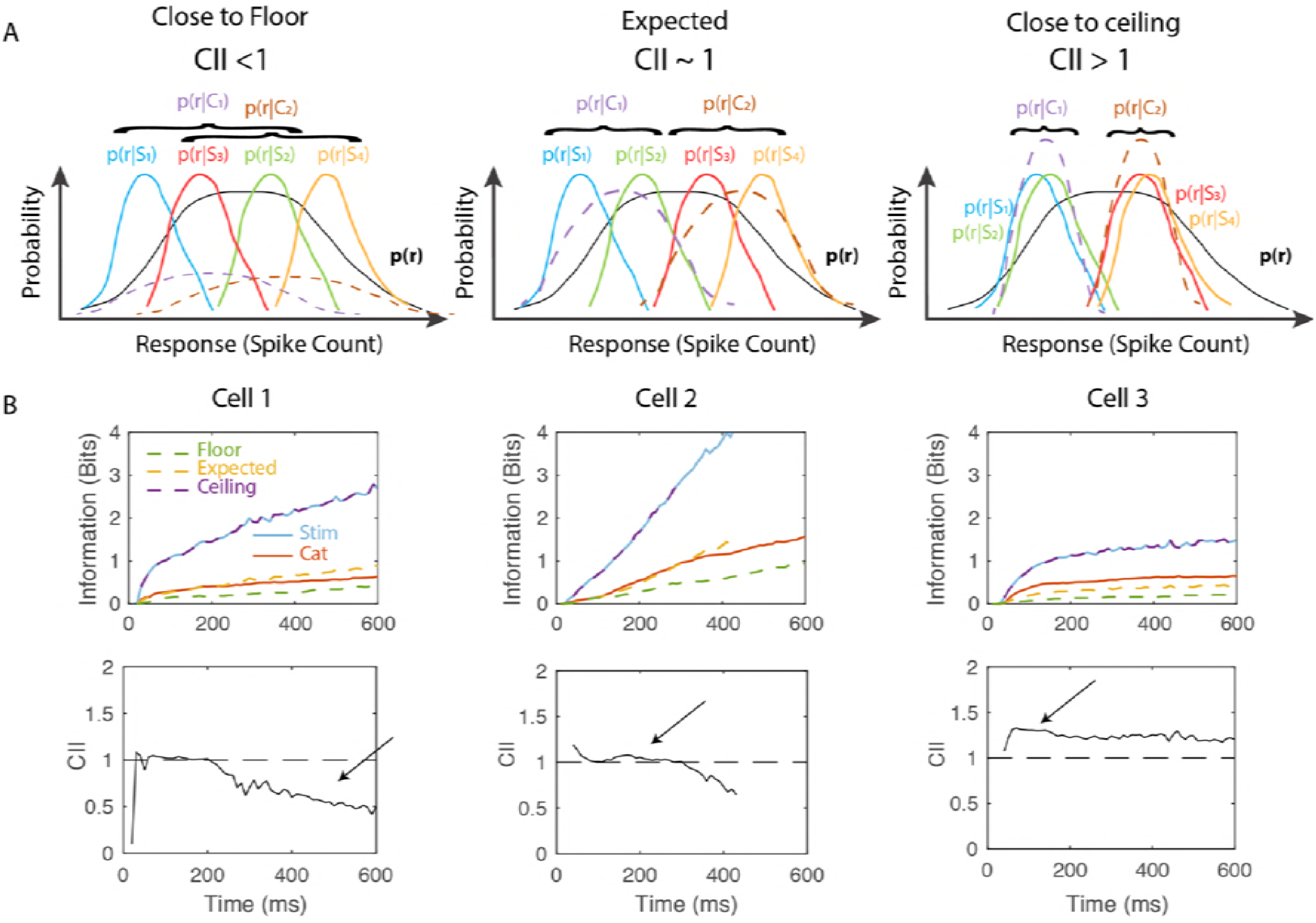
Cumulative Information and the Categorical Information Index. **A**. Schematic of hypothetical response probability distributions that yield different values of the Categorical Information Index (CII). CII is defined by comparing the cumulative categorical information to a floor (CII = 0), an expected value predicted from the stimulus information (CII=1) and a ceiling value (CII = 2) as described in the text and methods. Each plot represents the distributions of hypothetical neural responses for different stimuli (S1 to S4) that yield different values of CII. When stimuli are grouped randomly into categories (S_1_ and S_2_ belong to C_1_, while S_3_ and S_4_ belong to C_2_), CII is close to 0 (left panel). When stimulus categories preserve the order of the stimulus conditioned response distributions and these distributions are equally separated, effectively coding for both stimuli and categories, CII is equal to 1 (middle panel). When stimuli belonging to the same category have identical responses, the stimulus information is equal to the category information, the response being effectively invariant within categories and CII is equal to 2 (right panel). Note that the actual response probability distributions are multidimensional (one dimension for each time lag) with Poisson marginals. **B**. Example of the cumulative information for stimuli and call-type categories and CII for 3 different representative neurons. The blue and red lines are the cumulative information for stimuli and categories respectively. Estimates of the floor (dashed-green), expected (dashed-orange) and ceiling (dashed-purple, here overlapping with blue) values for the categorical cumulative information as described in A are given as well. The CII is shown as a solid line in the second row for each example neuron. Cell 1 has long periods of time with CII < 1, Cell 2 has long periods of time with CII ~ 1 and Cell has long periods of time with CII > 1 as shown by the arrows.

The plots in Figure 8 show the results of this analysis for the population both in absolute information units (left panel) and in the normalized units of CII. The thin colored lines correspond to CII curves for single neurons and they are colored according to the time average CII. The average CII over neurons (bold line on right panel) is very close to the shared value of 1 as one might expect if acoustical differences across stimuli drive neural responses in a linear fashion along some acoustical feature *and* call categories segregate perfectly along that same acoustical feature. The average CII is slightly (and significantly) above 1 between 120 and 260 ms showing some small degree of average invariance for call renditions within a call-type category during those times. Focusing, on the 25% of neurons with the highest CII, we found a peak at 175 ms. More significantly, perhaps, it is clear that there is a wide distribution of CII around the shared value of 1. However, this distribution includes many neurons that exhibit a high degree of invariance for call renditions within call-type categories as shown by the average CII for the top quartile (red solid line). For that top 25%, the absolute value of additional categorial cumulative information relative to the expected shared information value reaches a maximum of 0.16 bits at 320 ms (Figure 8A).

**Figure 8.**
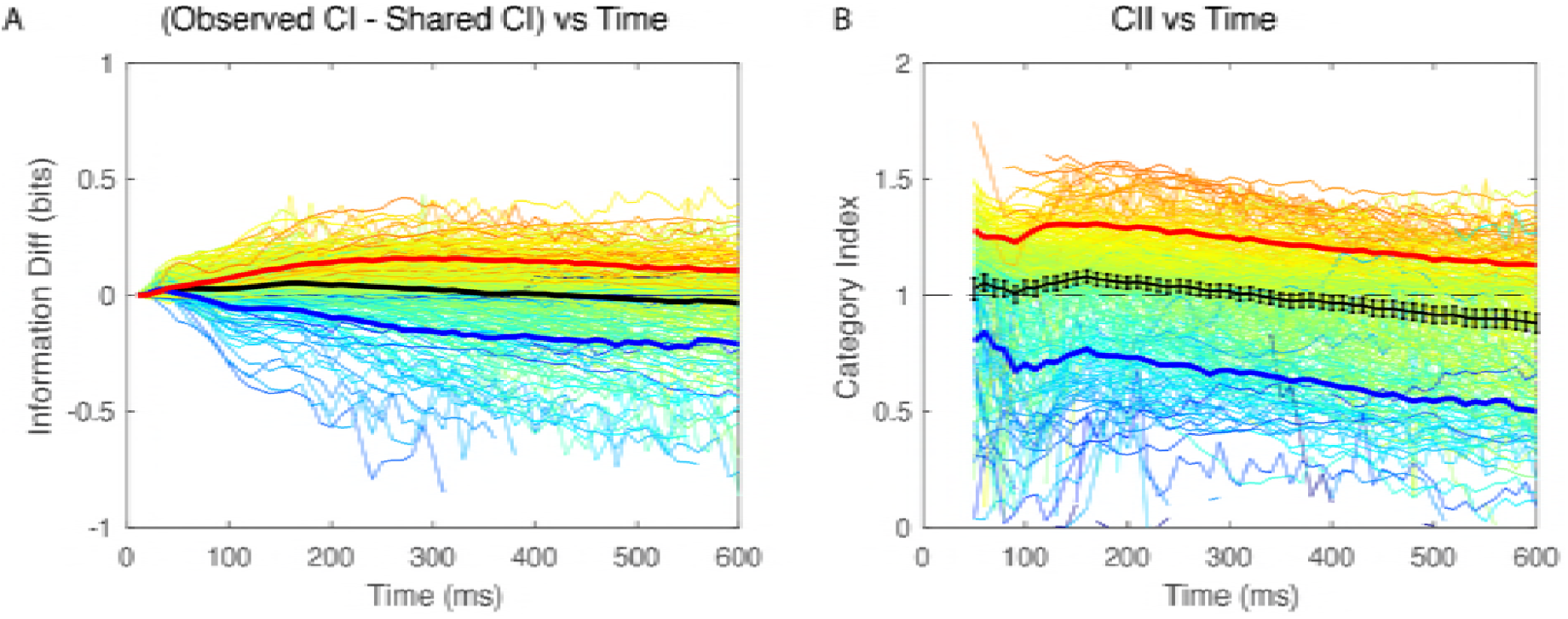
Categorical Cumulative Information. As explained in Fig. 7 and in the text, the cumulative information (CI) for call-type categories can be analyzed in terms of a floor or minimum value, a shared value and a ceiling or maximum value. **A**. Difference in the observed cumulative information in bits relative to the shared value. The bold black line is the average across all neurons (n=337). The thin colored lines correspond to individual neurons. Each line color corresponds to the time average Categorical Information Index (CII) of the neuron obtained between 50 and 300ms. The solid red line is the average obtained for the quartile of neurons with the highest CII and the blue solid line is average obtained for the quartile of neurons with the lowest CII. The same color code is used in B and in Figs. 9 and 10. **B**. Value of the Categorical Information Index as a function of time for all neurons in the population. Error bars on the average CII (solid bold black line) are 2 sem.

Do these high CII neurons exhibit other characteristic response properties? In the scatter plots of Figures 9 and 10, we examined the relationship between CII (color coded) and, respectively, time constants and relative level of the cumulative information. It can be seen from Figure 9, that neurons with high CII have relatively long stimulus time constants in comparison to their corresponding time constant observed for categorical information. This relationship is also significant for the entire population of neurons (Linear Regression explaining CII from *τ_cat_*/*τ_stim_*: Adj R^2^=0.11, F(1,213) = 28.5, p=2.37 10^−7^). As shown in Figure 10, neurons with high CII also have higher relative levels of categorical information in comparison to their relative values for stimulus information although this result is expected given our definition of CII (Linear Regression explaining CII from k_cat_(300)-k_stim_(300): Adj R^2^=0.76, F(1,213) = 675, p=5.5 10^−68^). Finally, one can also notice that neurons with high CII have low values of relative cumulative information (Linear Regression explaining CII from k_stim_(300): Coef = −0.84 Adj R^2^=0.28, F(1,213) = 84, p=3.96 10^−17^). This effect is caused by the correlation between invariance and selectivity as we have shown previously[40]: neurons that show the highest degrees of invariance also tend to respond to a small number of call-type categories and thus a small number of stimuli. As such, high invariance and high selectivity goes hand in hand with lower values of information.

**Figure 9.**
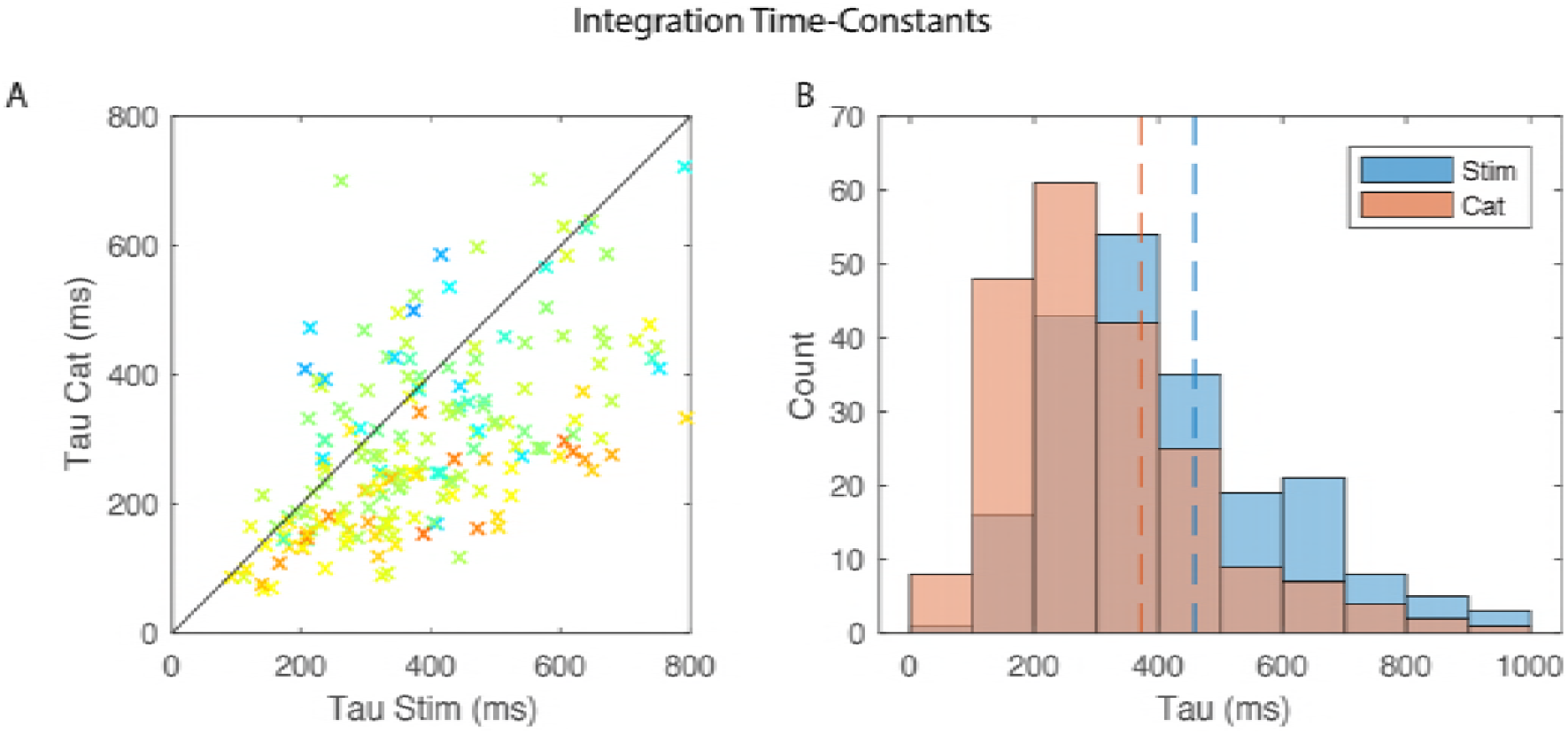
Time constants for Cumulative Information. For a large fraction of neurons (214/337 neurons), cumulative information for stimuli and call-type categories were well fitted with exponential curves. **A**. Scatter plot of the time constant of the exponential fit for stimuli (Stim – x axis) versus call categories (Sem – y axis). Single data points are colored according to the time averaged CII of each neuron (see Fig 8). **B**. Histogram of the two distributions of time constants. Dashed lines indicate the mean of each distribution (Stim = 459 ms; Cat = 372 ms).

**Figure 10.**
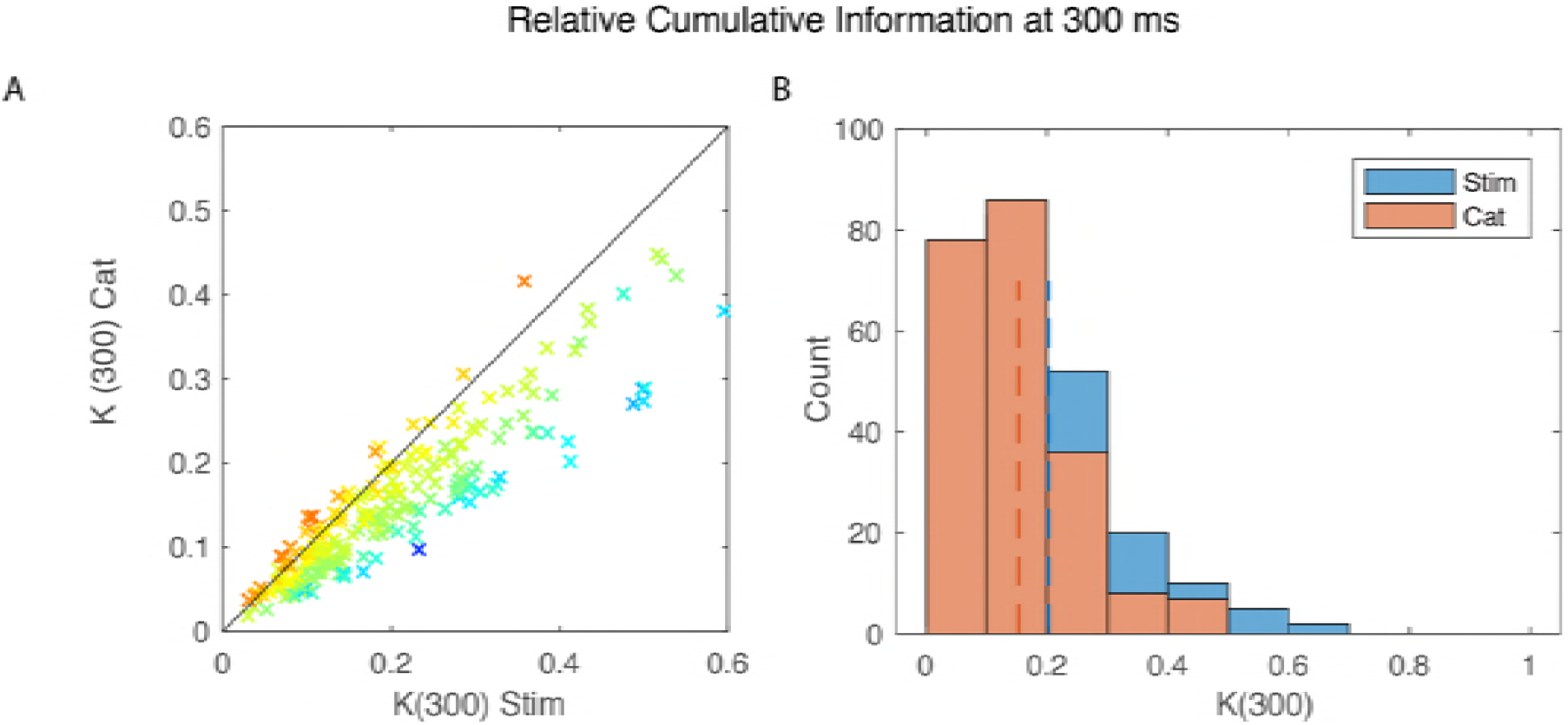
Distribution of Relative Cumulative Information estimated at 300 ms. For a large fraction of neurons (214/337 neurons), cumulative information for stimuli and call-type categories were well fitted with exponential curves. Here we used these fits, to estimate the cumulative information obtained at 300 ms relative to the maximum achievable information (log_2_(n_stim_) or log_2_(n_cat_)), k(300). k(300) is a number between 0 and 1 **A**. Scatter plot of k(300) for stimuli (Stim – x axis) versus categories (Cat – y axis). Single data points are colored according to the time average Categorical Information Index (see Figs 7–8). **B**. Histogram of the two distributions of k(300). Dashed lines indicate the mean of each distribution (Stim = 0.2; Cat = 0.15).

### Anatomical Organization

We examined whether neurons were organized in the avian auditory cortex based on their CII, cumulative information time constants (*τ*) and saturation levels (*k*). Most of our recording sites were identified histologically and could be assigned to avian cortical areas that had been segregated into the thalamic recipient area, L2; intermediate primary auditory regions, L1, L3, CML and L; and secondary areas, CMM and NCM [43–46]. We also obtained spatial x,y,z coordinates of the recording sites relative to the midline, the position along the rostral-caudal axis where the lamina pallio-subpallialis LPS) is the most dorsal, and the top of the brain (Fig. 11).

**Figure 11.**
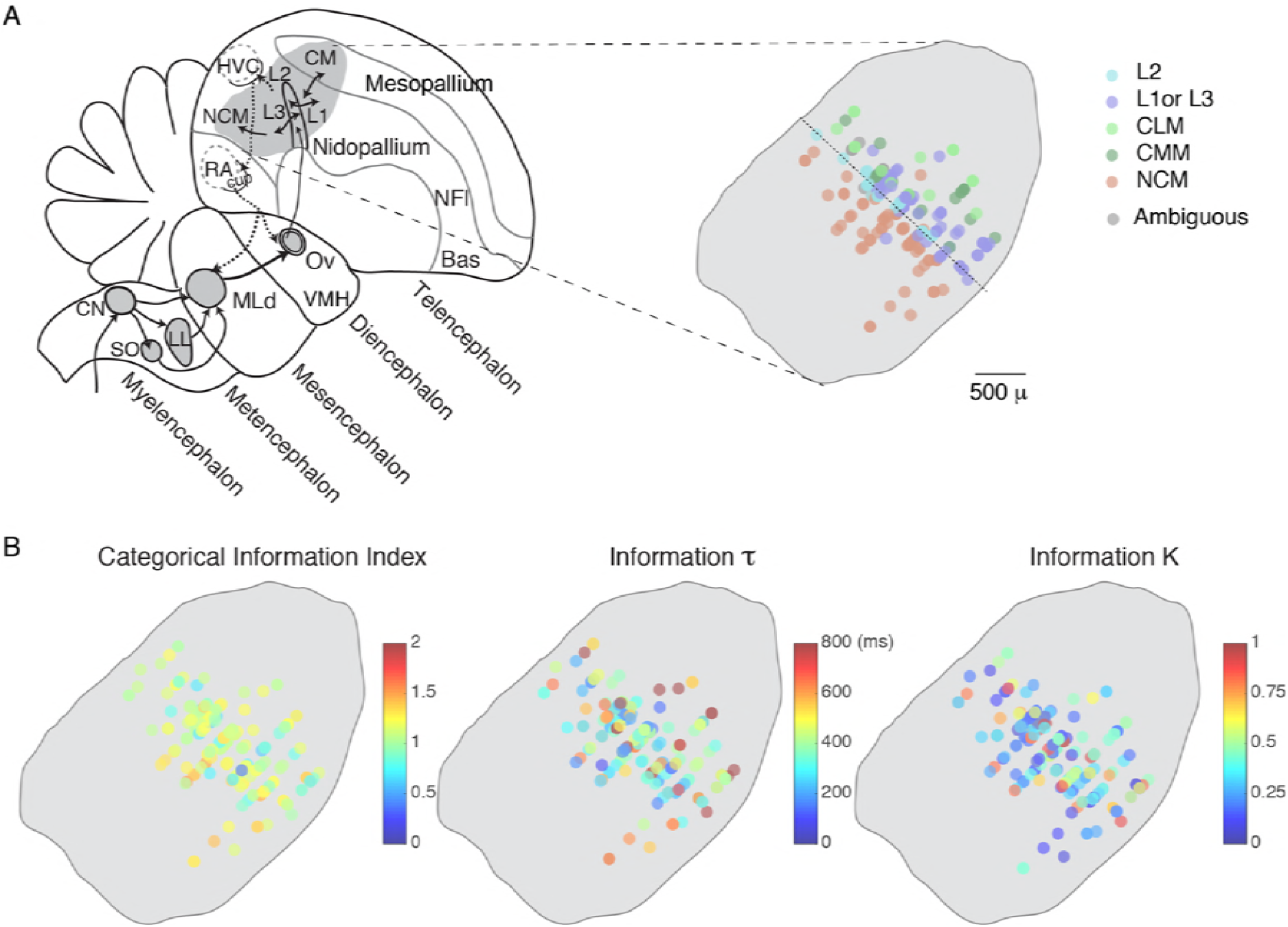
Anatomical Organization. **A**. The left panel is a schematic of the different processing stages in the ascending avian auditory system (grey areas) shown on a parasagittal view. Our recording sites were from the primary and secondary avian auditory cortical areas (the auditory pallium) and are shown on the right panel. The recording sites from different birds were aligned in the rostral-caudal dimension using the oblique rostro-caudal *long* axis of L2 (dotted line). Recording site positions also varied along the medial-lateral dimension (in and out of the page) and are all superimposed here. **B**. Functional anatomical maps of the Categorical Information Index, the time constant *τ* and saturation value *κ* of the cumulative information for stimuli. Each site is colored according to the value(s) of the variable being displayed. Transparency is used to reveal overlapping sites in this view as well as, potentially, combination of single units that were recorded from the same electrode and site.

In all regions, we found a range of CII and thus only weak anatomical trends across areas. Using the L2 dorsal-ventral oblique axis as reference point, CII was slightly higher as one moved rostrally or caudally to higher regions of auditory processing (Linear Regression: Adj R^2^= 0.03, F(3,194)=4.19, p=0.016). However, an ANOVA also suggested that both regions NCM and L2 had slightly higher mean CII (Adj R^2^=0.027, F(4,185) = 2.77, p=0.042). We also observed an increase in the time constant (τ) for the stimulus cumulative information that parallel the increase in CII as one moved away from the L2 axis (Linear Regression: Ajd R^2^=0.03, F(3,194)=4.79, p=0.0093). These significant anatomical trends were characterized by very small effect sizes. We also did not find any anatomical organization of saturation constants (*k*) for the cumulative information; neurons that had high levels of information could be found next to neurons with much lower levels and similar levels of average information were found in all regions.

## Discussion

We modeled responses that are observed in the auditory system as inhomogeneous Poisson processes in order to estimate the time-varying instantaneous and cumulative information for vocalizations used in communications. We showed that using Kernel Density Estimation for the time-varying firing rate and Monte Carlo with importance sampling for estimating probabilities, we were able to obtain accurate and bias-free estimates of these time-varying information values. This parametric approach is powerful because a relative small number of trials can be used to estimate the time-varying response and thus information values can be estimated in response to a relative large set of stimuli (here over 100 distinct vocalizations) given typical recording times. Poisson statistics were observed in the auditory cortical neurons recorded here when they were stimulated with short natural communication calls. More generally, the same procedures can also be used with spike trains statistics that can be parametrized with probability functions that depend only on a time varying rate such as inhomogeneous gamma or inhomogeneous inverse Gaussian [12, 47] and could also be extended to ensemble of neurons. Although Poisson statistics are often observed in neural data and will correctly fit any data set obtained from pooling responses across a large number of trials [48], it makes the strong assumption that the firing rate for a particular trial for a particular neuron depends only on time. Refractory period in spiking neurons and other correlations in single neurons or in an ensemble of neurons that are not phased locked to the stimulus (i.e. noise correlations) are common violations of this assumption. Taking noise correlations into account can increase stimulus decoding accuracies [23]. In cases where noise correlations are informative, the method proposed here would only yield lower bound estimates of information theoretic values. Alternatively, one should try the use of spike metrics measures [49] in combination with stimulus decoding approaches to then obtain measures of information from the confusion matrix of predicted versus actual stimuli [13, 40, 50, 51]. If spike metrics can be estimated accurately (with limited data), repeating the stimulus decoding procedure for progressively longer time windows would yield cumulative information values such as those calculated here but which could consider noise correlations.

Beyond our methodological contribution, the principal goal of this analysis was to characterize the neural code in higher auditory areas for communication calls. We found that, on average, auditory cortical neurons responded to these natural stimuli with time-varying firing rates that exhibited an onset and a sustained component of the response. Although in some situations the auditory cortex appears to respond only transiently [26, 52], our data supports findings from mammalian species that have showed strong sustained responses when neurons are driven by preferred stimuli [21]. Similarly, in the human superior temporal gyrus, the sustained responses to speech have been shown to be more informative for decoding speech phonemes [53]. Natural sounds in general have also been shown to be particularly efficient at driving auditory areas [12, 54–59] and thus the presence of informative sustained responses is not surprising. Indeed, in some of our most selective neurons, such as the neuron in Figure 5, the onset response is missing and only the sustained response is observed. More generally, and on average, we found that both the transient and sustained response had information about the stimulus identity *and* that the information in these two response phases was not redundant: the stimulus space of natural vocalizations is very large and although the initial response provides some clue as to the nature of the vocalization, new and additional information is observed in the sustained response. On the one hand, one might argue that, in natural vocalizations, the sound itself is changing with time. These stimulus changes occur both within calls that are made of composite notes or in call bouts and song motifs made of multiple syllables. Sustained responses in such cases could be interpreted as multiple onset responses (but with decreasing amplitude). On the other hand, from a behavioral perspective, these communication calls correspond to a single auditory object: a message from a particular individual about a particular state. From either perspective, these observations and analyses illustrate the importance of using behaviorally relevant stimuli when analyzing the nature of the neural code.

The information about stimulus identity was shown to be approximately equal in bits/s in the onset and sustained response. Given that the average onset response (in spikes/s) is greater that the sustained response, one might conclude that the onset coding (in bit/spikes) is not as efficient. Although, this is true from the point of view of a single neuron, it is almost certainly not true when ensemble of neurons is considered; the relative timing of the first spike has been shown both in audition [26] and in other sensory modalities [25, 27] to be highly informative. Moreover, in addition to stimulus identity other stimulus features are also encoded in neural responses; the timing of the stimulus is clearly marked by the onset response or transient response [22, 53, 60, 61] but other stimulus attributes such as the location, fundamental and loudness/distance of the sound source are also processed in auditory cortex [62–64]. It is therefore very likely that the onset response contains information about stimulus features other than stimulus identity or category and, potentially to a greater extent than in the sustained response (e.g. relative timing of onset). We note, however, that the opposite was found for the neural discrimination of two vowel sounds in the ferret auditory cortex where the onset response or early response was the most informative for decoding vowel identity relative to other attributes [64].

Our IT analysis also revealed the importance of stimulus locked spike patterns even in relatively short neural responses: in just 100 ms, the mean rate captured only a quarter of the information present when time-varying firing rates are considered. Thus, our analyses provide additional evidence that spiking patterns carry a significant amount of stimulus information and should therefore not be ignored in the analysis of neural responses [65]. The spiking precision analyzed here was relatively coarse (10 ms windows) and matched to the time scales of the relevant dynamics in the stimulus: although zebra finch calls are much longer than 10 ms, they are complex sounds with fast spectro-temporal features [39]. Therefore, although the neural code observed here uses fast varying spike rates, it cannot be labelled as a temporal code. In a rigorous definition of a temporal code, temporal information in spike patterns must code stimulus attributes other than the stimulus dynamics [66].

The cumulative information for stimulus identity or for categories increased for sustained periods of time before saturating. These saturating curves were well fitted with exponentials and yielded relatively long-time constants of approximately 460 ms for stimuli identity and 370 ms for call categories. These *information* time constants are long in comparison to the integration times that are usually found for auditory cortical neurons; the spectro-temporal receptive fields of avian auditory neurons rarely extend beyond 50 ms [67–70] although adaptive responses on longer time scales have also been described [71–73]. Information time constants depend on the integration and adaptation time constants of neurons but *also* on the stimulus dynamics: although stimulus identity and stimulus category are fixed in time, the sound itself has time varying features that can provide additional information as time goes on. It is the triple combination of the dynamics of the natural stimulus statistics, the neuronal integration time and the neuronal signal to noise levels that are going to affect the information time constant. Natural sounds and in particular communication calls are informative objects not only because of their spectral structure but also their rich dynamical structure. The importance of time in the neural code used in the auditory system has been emphasized multiple times [74–78] and our cumulative information analysis further stresses the importance of using natural sounds or synthetic sounds that carefully match natural dynamics when probing auditory neural encoding. Ultimately, it is an information time constant that includes stimulus dynamics that should be compared to behavioral responses [18]. A behavioral assay of reaction time for all the calls in the zebra finch repertoire has not been performed but the values of a few hundred of ms correspond to the shortest time intervals between call and call back in anti-phonal calling in paired zebra finches [79]. Although, faster times could be obtained in reaction to any sounds (e.g. startle or orientation), the processing of stimulus identity to extract the information on who is calling and what is being said might require these longer processing times.

Finally, we quantified the fraction of the information about stimulus identity that could be used for extracting the call-type category. We used that analysis to characterize neurons not only in terms of their absolute coding capacities for call-type categories but also in terms of their invariance in their response to different stimuli belonging to the same call-type category. We found that, on average, information for categories is very close to what one would expect if neural representation for stimulus identity is also segregated along call-type categories. However, we also found neurons that had a high degree of invariance and could therefore be classified as categorical. On the one hand, these categorical and invariant neurons had even longer time constants for cumulative stimulus information suggesting that they could be higher in the auditory processing stream. On the other hand, we did not find a separate population of invariant neurons across all of our recordings of neurons informative for categories: the distribution of our categorical index was unimodal. Moreover, we found only weak anatomical correlations with very small increases in the Categorical Information Index along an anatomical axis corresponding to lower vs higher auditory areas. Higher avian auditory cortical areas have been associated with more complex spectro-temporal receptive fields [67, 70, 80], increase robustness to noise [81–83] and more specificity for processing natural sounds [12, 84]; but, it is also striking to see that in all those analyses as well as those that focused on single avian auditory cortical areas [85] there is a high degree of heterogeneity in the neural responses within each area. Here, we also found that neurons with high Categorical Information Indices could be found anatomically next to neurons with low Categorical Information Indices. Some of this functional heterogeneity could be associated with different cell types [69, 84] and a better understanding of the micro-circuitry in the avian auditory cortical areas is needed [30]. It is interesting to note, however, that this mixing of low level and high-level response properties is not unique to the avian auditory system as similar heterogeneity has been found in the mouse [86] and ferret auditory cortex [87, 88]. Contrary to the visual system, the auditory system might preserve a higher mixture of low-level and high-level sensory responses properties at multiple stages of processing including the higher auditory areas involved in auditory object recognition. If this is true and universally found in vertebrates, it might be a necessary property of the computations needed for auditory object recognition, potentially related to the fact that complexity in auditory signals is in time-varying spectral patterns that quickly disappear; the fleeting nature of sounds could prevent higher processing stages from subsequently accessing lower-level representations for additional information.

## Methods

### Animals and Stimuli

Four male and two female adult zebra finches (*Taeniopygia guttata*) were used for the electrophysiological experiments. The birds were bred and raised in family cages until they reached adulthood, and then maintained in uni-sex groups. Although birds could only freely interact with their cage-mates, all cages were in the same room allowing for visual and acoustical interactions between all birds in the colony. All birds were given seeds, water, grid and nest material ad libitum and were supplemented with eggs, lettuce and bath once a week. All animal procedures were approved by the Animal Care and Use Committee of the University of California Berkeley and were in accordance with the NIH guidelines regarding the care and use of animals for experimental procedures.

Vocalizations used as stimuli during neurophysiological experiments were recorded from 15 adult birds and 8 chicks (20-30 days old). The vocalization bank obtained contains 486 vocalizations that included for each bird most of the calls in the Zebra finch repertoire: 7 call-types in adults and 2 in chicks. The adult calls included the following affiliative calls: Song (So), Distance Call (DC), Tet call (Te) and Nest Call (Ne); and the following non-affiliative calls: Wsst or aggressive call (Ws), the Distress Call (Di) and one of the two alarm calls, the Thuk (Th). The juvenile calls included the Begging call (Be) and the Long Tonal Call (LT). Additional information about these stimuli and their behavioral meanings can be found in [39, 40, 89].

For the neurophysiological experiments, a new subset of the vocalization bank was used at each electrophysiological recording site. This subset was made from a representative number of vocalizations from the repertoire of individuals: three adult females, three adult males, two female chicks and two male chicks. From each individual caller, we randomly chose 3 call bouts from each category or fewer if fewer than 3 call bouts were obtained for that particular call-type and individual. The maximum number of stimuli that could be selected in that procedure was therefore 3×7×3 (adult males x repertoire x renditions) + 3×6×3 (adult females x repertoire x renditions) + 4×2×3 (juveniles x repertoire x renditions) = 141. Fewer stimuli were used when we had fewer renditions than 3 for a particular bird or when the signal from a single unit was lost before the end of the recording. The average number of stimuli played for each single unit was 114 (sd = 22, min = 34, max = 123). Ten trials were acquired for each stimulus with a few exceptions (min=9, max = 11). Sounds were broadcasted in a random order using an RX8 processor (TDT System III, sample frequency 24414.0625 Hz) connected to a speaker (PCxt352, Blaupunkt, IL, USA) facing the bird at approximately 40cm. The sound level was calibrated on song stimuli to obtain playbacks at 75dB SPL measured at the bird’s location using a sound meter (Digital Sound Level Meter, RadioShack).

### Neurophysiological and Histological Procedures

Extra-cellular electrophysiological recordings were performed in 6 urethane anesthetized adult zebra finches. The birds were placed in a sound-attenuated chamber (Acoustic Systems, MSR West, Louisville, CO, USA) and sound presentation and neural recording were performed using custom code written in TDT software language and TDT hardware (TDT System III). Sounds were broadcasted in a random order as described above. Neural responses were recorded using the signal of two (5 subjects) or one (1 subject) 16-tungsten electrode arrays, band-pass filtered between 300Hz and 5kHz and collected by an RZ5-2 processor (TDT System III, sample frequency 24414.0625 Hz). The electrode arrays consisted of two rows of 8 electrodes with row separation of 500 mm and inter-electrode separation within row of 250 mm. Electrode impedances were approximately 2 MOhms. When two electrode arrays were used, they were placed each in one hemisphere. Spike arrival times and spike shapes of multiple units were obtained by voltage threshold. The level of the threshold was set automatically by the TDT software using the variance of the voltage trace in absence of any stimuli. Electrodes were progressively lowered and neural responses were collected as soon as auditory responses to song, white noise, Distance call or limited modulation noise could be identified on half of the electrodes in each hemisphere (the stimuli used to identify auditory neurons were different from the stimuli used in the analysis). Several recording sites were randomly selected by progressively deepening the penetration of the electrodes and ensuring at least 100 μm between two sites. On average 4.2±2 sites (mean ± sd) were recorded per bird and per hemisphere at a depth ranging from 400 μm to 2550 μm.

After the last recording site, the subject was euthanized by overdose of isoflurane and transcardially perfused. Coronal slices of 20μm obtained with a cryostat were then alternatively stained with Nissl staining or simply mounted in Fluoroshield medium (F-6057, Fluoroshield with DAPI, Sigma-Aldrich). While Fluoroshield slices were used to localize electrode tracks, Nissl stained slices were used to identify the position of the 6 auditory areas investigated here: the three regions of Field L (L1, L2 and L3), 2 regions of Mesopallium Caudale (CM): Mesopallium Caudomediale (CMM) and Mesopallium Caudolaterale (CLM); and Nidopallium Caudomediale (NCM). By aligning pictures, we were able to anatomically localize most of the recording sites and calculate the approximate coordinates of these sites. Since we could not localize the Y-sinus on slices, we used the position of the Lamina Pallio-Subpallialis (LPS) peak as the reference point for the rostro-caudal axis in all subjects. The surface of the brain and the midline were the reference for respectively the dorsal-ventral axis and the medial-lateral axis. The approximate coordinates of units were used to build 3-D reconstructions of all single unit positions in an hypothetic brain.

Single unit isolation was performed off-line using custom software that used a combination of supervised and unsupervised clustering algorithms. These clustering algorithms used the spike-snippets shape as described by a PCA. Sorted units where declared to be single units based on spike shape reliability across snippets. The spike shape reliability measure was a signal to noise ratio (SNR) where the signal was the difference between the maximum and the minimum of the average snippet and the noise was the standard deviation of this measure across all snippets. Single units in our data set have an SNR > 5. Additional details on these experimental procedures can be found in [40].

### Data Analysis: Time-varying Spike Rate Estimation

The data of neural responses from 404 out of 914 isolated single units were used in this study. The 404 were selected based on a prior analysis that showed that this subset of units were not only auditory but also contained information about call-types, in the sense that call-types could be decoded above chance level from neural responses (see [40]). Here, we analyzed the neural response in the first 600ms after stimulus onset. For each stimulus, the 9 to 11 raw spike patterns of 600ms, sampled at 10kHz, were combined to obtain the corresponding time varying spike rate (sample frequency set at 1kHz) by applying a locally adaptive kernel bandwidth optimization method [41]. In cases where the neuron did not respond to any of the presentations of the stimulus or responded only once over all presentations, the rate was estimated as being constant for the 600ms duration of the neural response. For those two unresponsive cases, the rate was set to be 1/(2*Ntrials*NTimes) in the absence of any spike or 1/(Ntrials* NTimes) in case of one spike, with Ntrials the number of stimulus presentations (9-11) and NTimes the number of time points at which the rate was estimated (here 600, for a 600ms neural response section with a sampling rate set at 1kHz).

Calculations of cumulative information requires the estimation of very large distributions which sizes grow exponentially with the number of time points investigated. To investigate cumulative information values up to 600ms after stimulus onset, time-varying rates were sampled at 10ms (Nyquist limit frequency of 50Hz). To estimate, the amount of information potentially lost by this low-pass filtering, we estimated an information value based on coherence analysis of the signal to noise ratio in the raw spike train. The coherence between a single spike train (R) and the actual time-varying mean response (A) 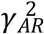 can be derived from the coherence between the peristimulus time histogram (PSTH) obtained from half of the trials and the PSTH obtained from the other half [90].

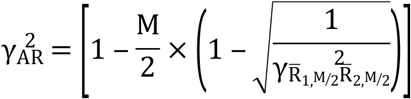

where M the total number of trials (presentations of the stimuli) and 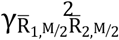 the coherence between the two PSTHs calculated on half of the trials. The coherence between two responses is a function of frequency (ω). An estimate of the mutual information (in bits per second) between R and A responses can then be obtained by integrating over all frequencies [4, 90]:

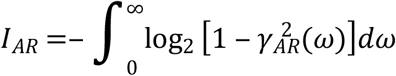

For each unit, we estimated the percentage of information preserved as the ratio between *I_AR_* calculated up to 50Hz and *I_AR_* calculated over all frequencies. Over all units, 96.7% ±6.9% (mean ± SD) of information was conserved by a lowpass filtering at 50Hz and only 88 out of 404 cells had information losses greater than 5%. For each unit, we also calculated the proportion of cumulative power across frequencies in the time varying spike rate estimation obtained with the KDE before low-pass filtering and down-sampling (averaged periodogram in overlapping 200 ms Hanning windows and 1 kHz sampling rate). The cumulative sum of the power was calculated across frequencies and normalized by the maximum power value to obtain the proportion of cumulative power. On average across units, the cumulative power reached 98.8%±1.6% (mean ± SD) at 50Hz, further validating our choice of the temporal resolution (Supp Fig. 1).

### Information Theoretic Calculations

As described in the results, the time-varying *instantaneous* mutual information between the stimulus S and the response *Y_t_* can be written as a difference in Shannon entropies:

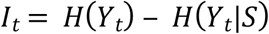

and the *cumulative* mutual information for neural responses that are discretized into time intervals is given by: *CI_t_* = *H*(*Y_t_, y*_*t*−1_*Y*_*t*−2_,…,*Y*_0_) – *H*(*Y_t_,y*_*t*−1_*Y*_*t*−2_,…,*Y*_0_|*S*) In the present paper we calculated 4 different types of information: the stimulus instantaneous information, the categorical instantaneous information, the stimulus cumulative information and the categorical cumulative information. Stimulus instantaneous and cumulative information were calculated for all 404 units, while categorical instantaneous and cumulative information were calculated on a restricted set of 337 neurons that presented at least one time point with a significant value of stimulus cumulative information (significant threshold set as 3 times the local error, see below for error calculations). While we verified our assumptions (minimal information loss with spike rate binning, Poisson distributions of spike counts and maximum value of spike counts) on the full set of 404 units, the population analysis of time-varying information presented in the results section only include the relevant dataset of 337 neurons.

The custom Matlab code used to calculate time-varying information values is available at https://github.com/julieelie/PoissonTimeVaryingInfo along with a tutorial on how to use the core functions.

### The Instantaneous Information

The conditional response entropy and the response entropy are obtained from the distribution of the conditional probability of neural responses given the stimulus, *p*(*y_t_*|*s*), and the distribution of probability of each stimulus *p*(*s_i_*):

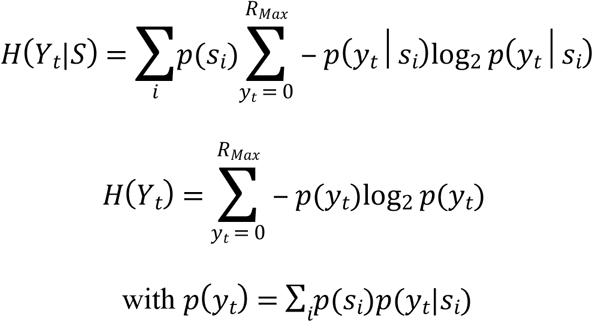

We modeled the distribution of neural responses to a given stimulus *Si* as an inhomogeneous Poisson process. The conditional probability of response (spike count) given the stimulus is then:

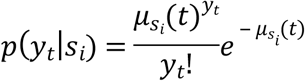

And the local entropy is:

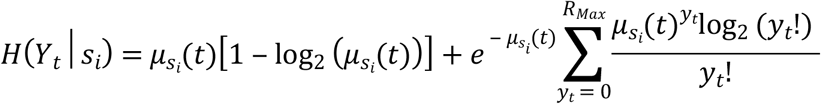

Because, in our data, the probability of response is very small for high values of *y_t_*, calculations of entropies were bounded for *y_t_* between zero and *R_Max_*. Here we set *R_max_* = 20 which corresponds to a rate of 2 spike/ms in the 10 ms analysis windows. The maximum rate observed in all of our neurons across all time bins was 0.8 spike/ms (Sup. Fig. 2). Note that this very high firing rate (800 Hz) was only observed only once; that is in one 10 ms time windows across all neurons (404) and all stimuli (114*60= 6840) or with a p=1/ 2763360. This is clearly the very upper limit of a distribution with a long tail. Our time-varying rates were well below that upper bound but we verified that the cumulative probability up to *R_max_* = 20 was numerically identical to 1 before estimating entropies. As described above (see time-varying spike rate estimation), we also enforced a lower bound for *μ_s_i__*.(*t*) of 1/20.

The total conditional entropy at time *t* is:

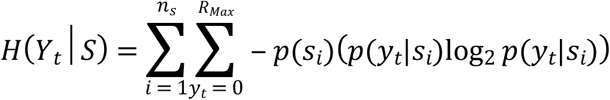

where *p*(*S_i_*) is the probability of observing stimulus *S_i_* and *n_s_* is the number of stimuli sampled: it is the average of the local entropies obtained for the conditional probability of response for each stimulus *s_i_*. Assuming that our sample is representative of stimuli encountered, each stimulus is equally probable, 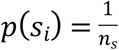 or:

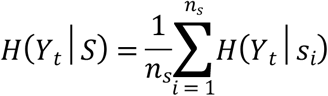

Alternatively, one can assume that each call category is equally probable. If *k_i_* is the number of stimuli in the particular call category *c* to which *s_i_* belongs and *n_c_* is the number of categories, then:

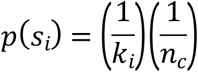

for *s_i_* ∈ *c*. Then:

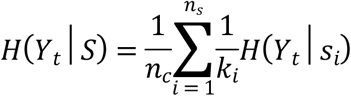

The unconditional probability (i.e. across all stimuli) of a response at time *t* is not Poisson but is given by a weighted sum of Poisson with different mean rates:

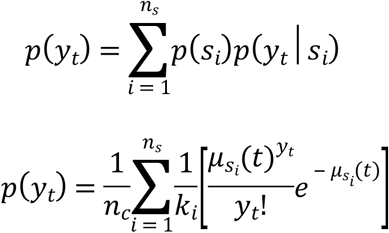

The response entropy is obtained from these unconditional probabilities:

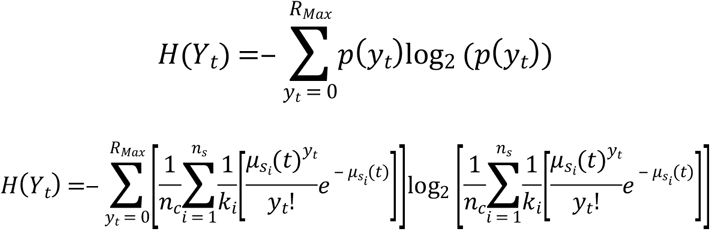

### The Cumulative Information

The conditional probability of a time varying response is the joint probability of observing (*y_t_,y*_*t*−1_,*y*_*t*−2_,…) given *s_i_*. Given our Poisson assumption, the conditional probability of response at *t* is independent of the conditional response at previous times.

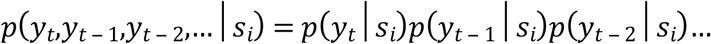

We can show that the conditional entropy of the joint responses is the sum of the individual entropies:

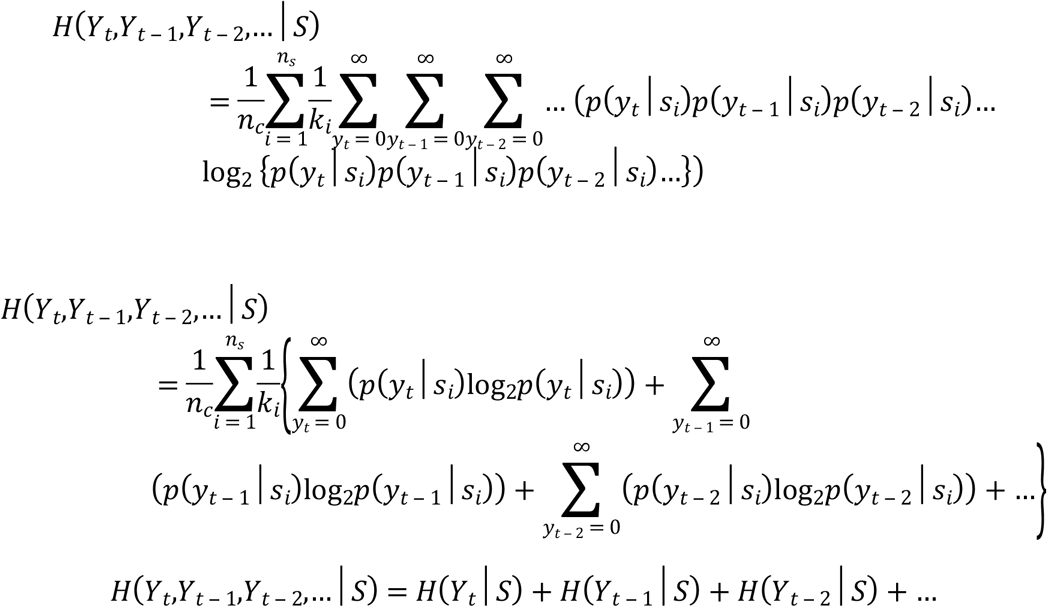

The probability of the time varying response is the joint probability of observing (*y_t_, y*_*t*−1_*y*_*t*−2_,…). This joint probability cannot be expressed as the product of the probabilities at different times because these are not independent. The joint unconditional probability distribution is:

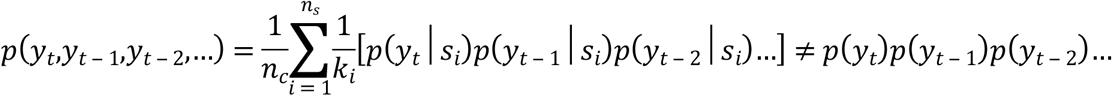

This joint probability distribution could be expressed as a product of probabilities by assuming that most of the interdependence can be calculated from the previous time point, as in the Markov chain assumption at a beginning of the time series at *t*=0. In all cases, the joint probability distribution can be written as:

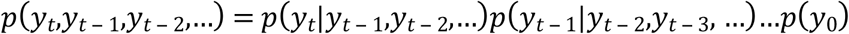

or when it is approximated by a first-order Markov chain:

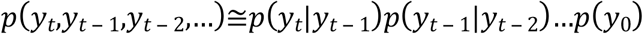

The conditional probability of *y_t_* given *y*_*t*−1_ is:

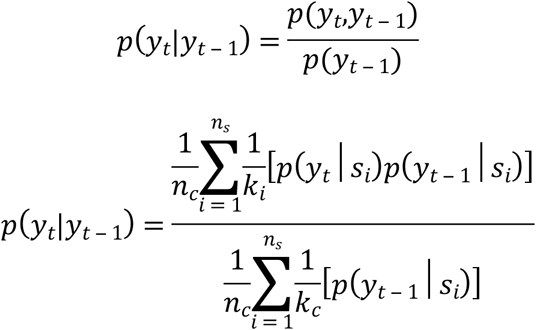

Note that this joint probability distribution can have very high dimensions. Assuming the number of spikes *y_t_* ∈ [0, 19], the number of probabilities that must be estimated is 20^*nt*^ where *nt* is the number of windows in time. For example, calculating all the probability of all the outcomes for 10 windows (100ms) requires 20^10^~10^13^ calculations. On the other hand, the Markov chain approximation only requires the estimation of all pair-wise joint probability distributions: for 10 windows and 20 outcomes, the number of calculations is (10)(20^2^)~4000.

The response entropy is then calculated from the joint probability distribution:

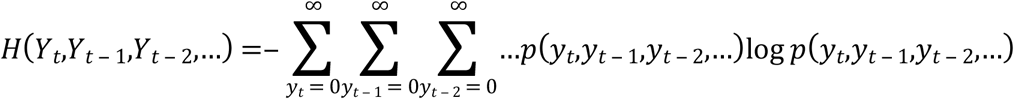

To estimate this response entropy, we investigated various approaches: a time-running cumulative information, the Markov chain approximation and Monte Carlo with importance sampling. Monte Carlo with importance sampling gave the best results and was therefore used in our analyses. We briefly describe the three approaches. The time-running cumulative information consisted in calculating the full cumulative information (using all possible spike events) but only for a fixed number of successive time windows. We estimated that we could easily calculate all possible probabilities for 4 time-windows, corresponding to a 40 ms history. This approach gave the best estimate of the information in 40 ms windows but, in our system, grossly underestimated the cumulative information: some of the information in successive 40 ms is clearly independent (Sup. Fig. 3).

The second approximation was based on Markov chain of variable orders up to 4 (also 40 ms). With this approximation, we overestimated the cumulative information: the correlation time of the time-varying rates for different stimuli is clearly also greater than 40 ms. Using the first order Markov chain approximation:

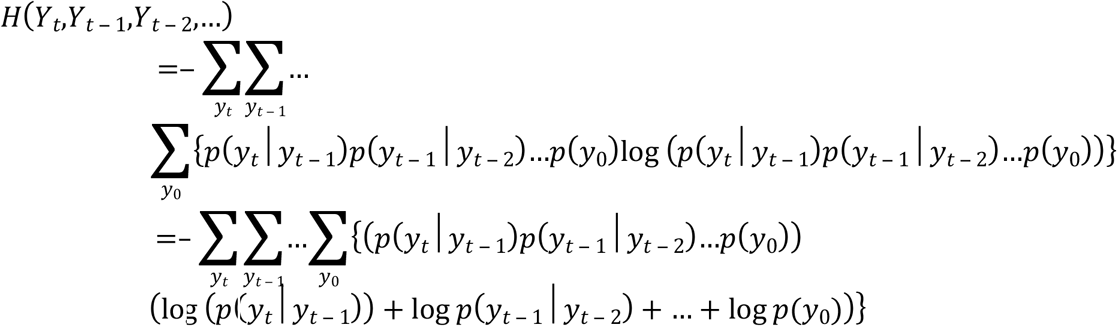

Expanding that sum, the last term is:

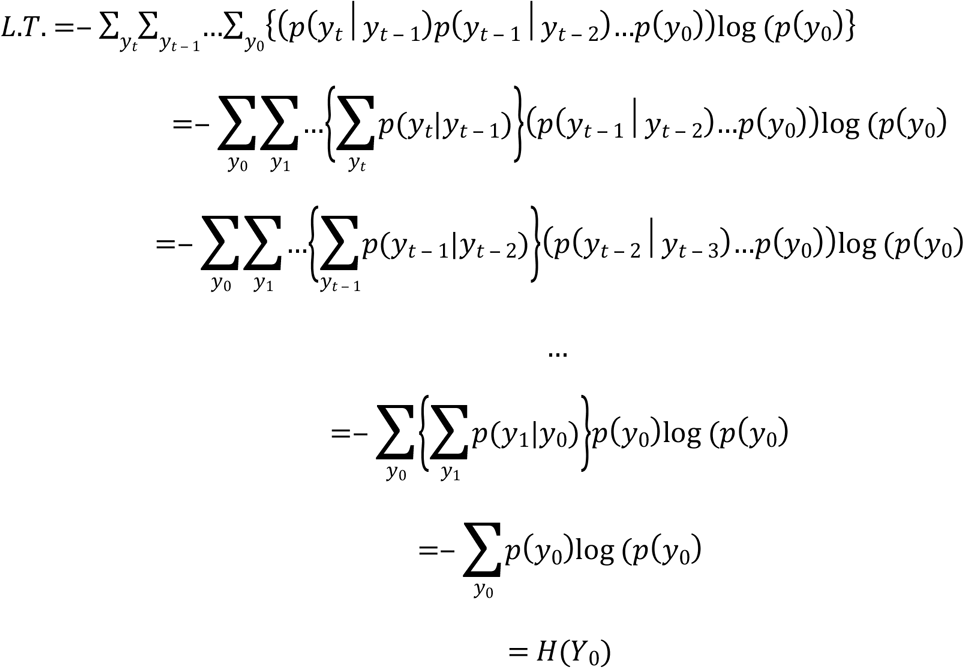

The second to last term is:

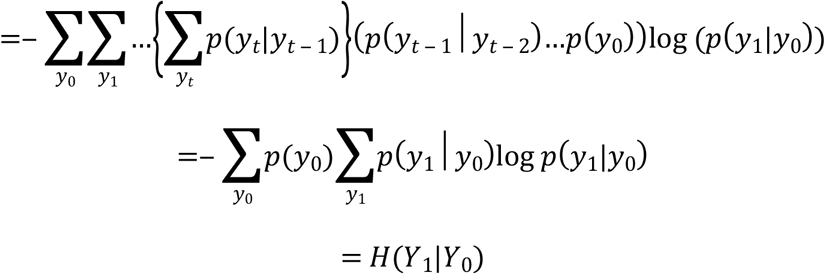

The third to last term is:

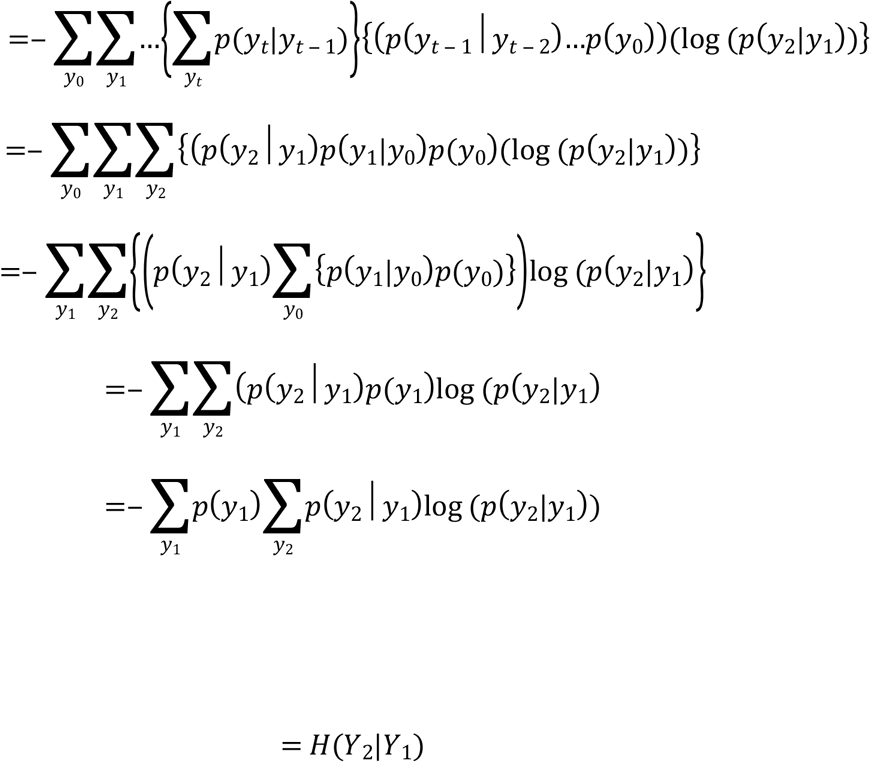

and, similarly, for all the other terms.

Thus, the response entropy using the Markov chain approximation is:

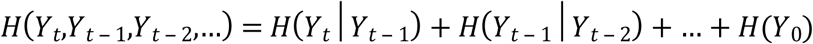

with *H*(*Y_t_*|*Y*_*t*−1_) =− Σ_*y*_*t*−1__[*p*(*y*_*t*−_)Σ_*y_t_p*_(*y_t_*|*y*_*t*−1_))]log_2_(*p*(*y_t_*|*y*_*t*−1_))]

This approximation can be extended using estimates at two previous time points, *etc*.:

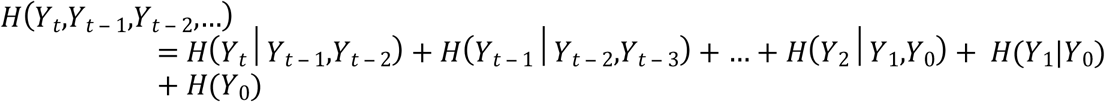

where the conditional probability based on two prior measurements is:

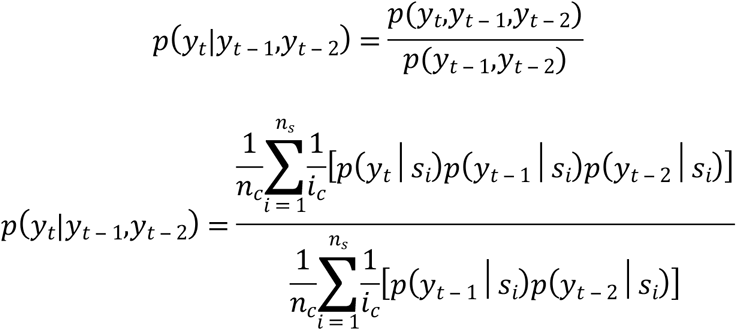

where *H*(*Y_t_*|*Y*_*t*−1_,*y*_*t*−2_)=−Σ_*y*_*t*−1__

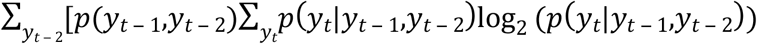

Finally, we estimated cumulative information using Monte Carlo with importance sampling. In Monte Carlo in conjunction with importance sampling, *N_i_* samples or, here, time varying responses 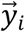, are taken from a proposal distribution, 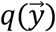. The actual probability, 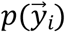, is calculated exactly at those samples and an estimate of the expected value of the measure of interest (here 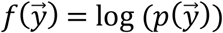) is obtained by the average of 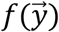 at the sample points, 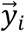, weighted by the likelihood ratio *p*/*q*:

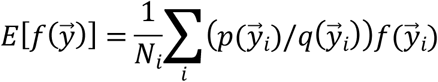

Our proposal distribution was based on the distribution at each time point and assuming independence across time:

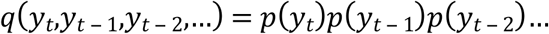

For each sample obtained from the proposal distribution q we calculated the actual probability value using:

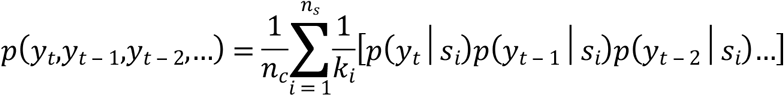

and used that probability value in the estimation of the entropy. Monte Carlo samples were chunked in groups of 100,000 samples and at each additional sample chunk, a bootstrapped/jackknife bias corrected mean and standard error were estimated (see below). Sampling stopped when the standard error was below 0.2 bits or at a max number of samples set to 5,000,000 samples. If the error at the maximum number of samples was greater than 0.6 bits, the estimation was deemed unreliable and the calculation was not performed for any successive time points.

### Bias Correction and Standard Error for Information Calculations

The small number of trials used to estimate spike rates is the source of bias and uncertainty in our calculation of information: a small number of stimulus presentations increase the probability of obtaining by chance estimated spike rate that are different between stimuli and so yields a positive bias on information calculations. We used a Jackknife procedure on the estimation of spike rate for each stimulus to correct for this positive bias. Jackknife kernel density estimate of the rate were obtained by applying the locally adaptive kernel bandwidth optimization method on the Ntrials possible sets of Ntrials-1 spike patterns of each stimulus (Ntrials being the number of stimulus presentation of a given stimulus). Moreover, uncertainty about information values comes from the sampling errors on spike rate and on the Monte Carlo estimation of joint spike rate probability distributions (cumulative information only). These errors on information calculations were estimated by bootstrapping the jackknife procedure:

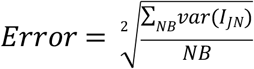with *NB* the number of boostrap (*NB* = 20), *I_JN_* the bias-corrected estimation of information obtained from the Jackknife procedure.

### Calculation of the Expected Value of Categorical Information Given the Stimulus Information

We computed a Categorical Information Index (CII) that compared the empirical categorical cumulative information for call-type categories, CCI, to three hypothetical values: a floor (*CCI_FLOOR_*), an expected value (*CCI_Exp_*) and a ceiling value (*CCI_Ceil_*). The floor value is the categorical cumulative information obtained from random categories. The expected value is the categorical cumulative information that would be achieved if the stimulus information was 1) equally distributed for each stimulus and 2) could be used for classifying stimuli into groups. Note that the second assumption is not necessarily true in the actual data because the categorical information is based on averaging the probabilities for stimulus from the same category and thus effectively averaging time-varying rates. If time-varying rates are not grouped by categories, then it is possible that two stimuli from two different categories are distinguishable based on their time-varying rate, but that, the average time-varying rates for the two categories are not distinguishable, or less so than expected from the average pair-wise distances. The ceiling value corresponds to the case where all the cumulative information about stimuli is used for the categorization and none to discriminate stimuli belonging to the same category: *CCI_Ceil_*. The CII is a number between 0 and 2 that is then calculated as:

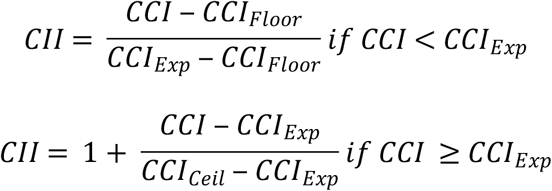

The following three steps were taken to calculate the expected categorical information (*CCI_Exp_*) from the stimulus cumulative information. First, the stimulus mutual information, *mi*, was expressed as the conditional probability of correct stimulus decoding, *p*, for any given stimulus (and assumed to be equal for all stimuli). Given a confusion matrix of size nxn obtained from a decoder for *n* stimuli, with *p* as the conditional probability given a stimulus of correct decoding (diagonal terms) and thus *(1-p)/(n-1)* as the conditional probability of error (off-diagonal terms), the mutual information is equal to:

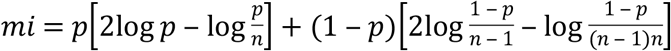

The above equation was inverted numerically to solve for *p* given *mi*. Second, a new matrix was generated by grouping rows and columns of joint probabilities (and not conditional) to form a confusion matrix for categories. The number of stimulus in each category was matched to the actual values in the neurophysiological data on a unit per unit basis. Third, the expected mutual information for categories was then estimated from this new confusion matrix by subtracting the total entropy obtain from the joint probabilities, from the sum of the entropies of the marginal distributions for the rows and columns:

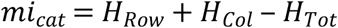

## Data availability

The custom Matlab code used to calculate time varying information values is available at https://github.com/julieelie/PoissonTimeVaryingInfo, along with a tutorial on modeled data.

## Acknowledgement

This research used the Savio computational cluster resource provided by the Berkeley Research Computing program at the University of California, Berkeley (supported by the UC Berkeley Chancellor, Vice Chancellor for Research, and Chief Information Officer).

## Supplemental Figures

**Supplemental Figure 1.**
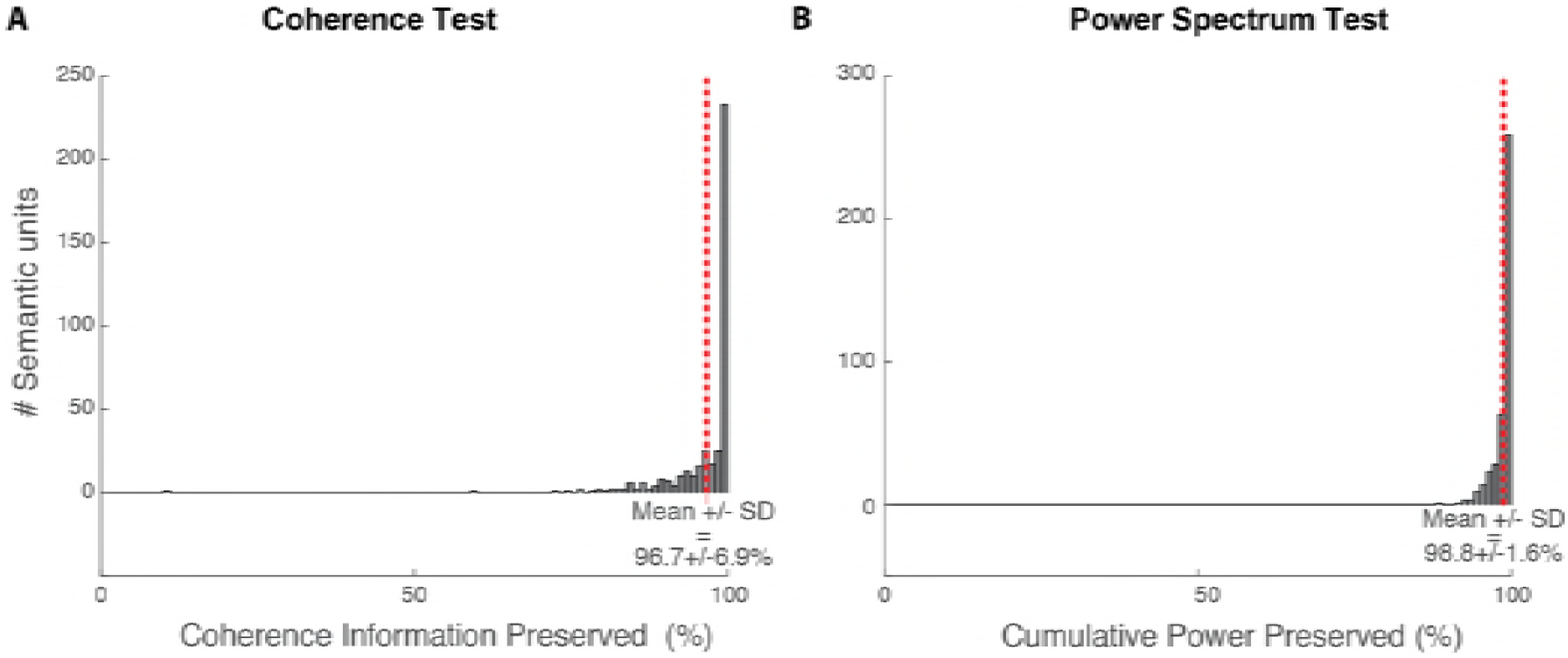
Tests for Temporal Resolution. We performed two tests to assess the potential information loss from sampling the time-varying rate at 50 Hz (10 ms bins). **A**. The Coherence Test is based on the coherence between individual spike trains. A measure of total coherence (Information Coherence) can be obtained by integrating over frequencies (see Methods). The Information Coherence obtained by integrating from 0 to 50 Hz can then be compared to the Information Coherence obtained for the entire frequency range of 0 to 500 Hz. The histogram shows the number of cells versus the fraction of Information Coherence in 0-50 Hz. **B**. The Power Spectrum Test is based on the power spectrum of the time-varying rates for each neuron obtained with the Kernel Density estimation (sample frequency: 1kHz). Just as for the Coherence Information, the fraction of the power between 0-50 Hz relative to the power between 0-500 Hz was estimated for all cells. The histogram shows the number of neurons as a function of that fraction.

**Supplemental Figure 2.**
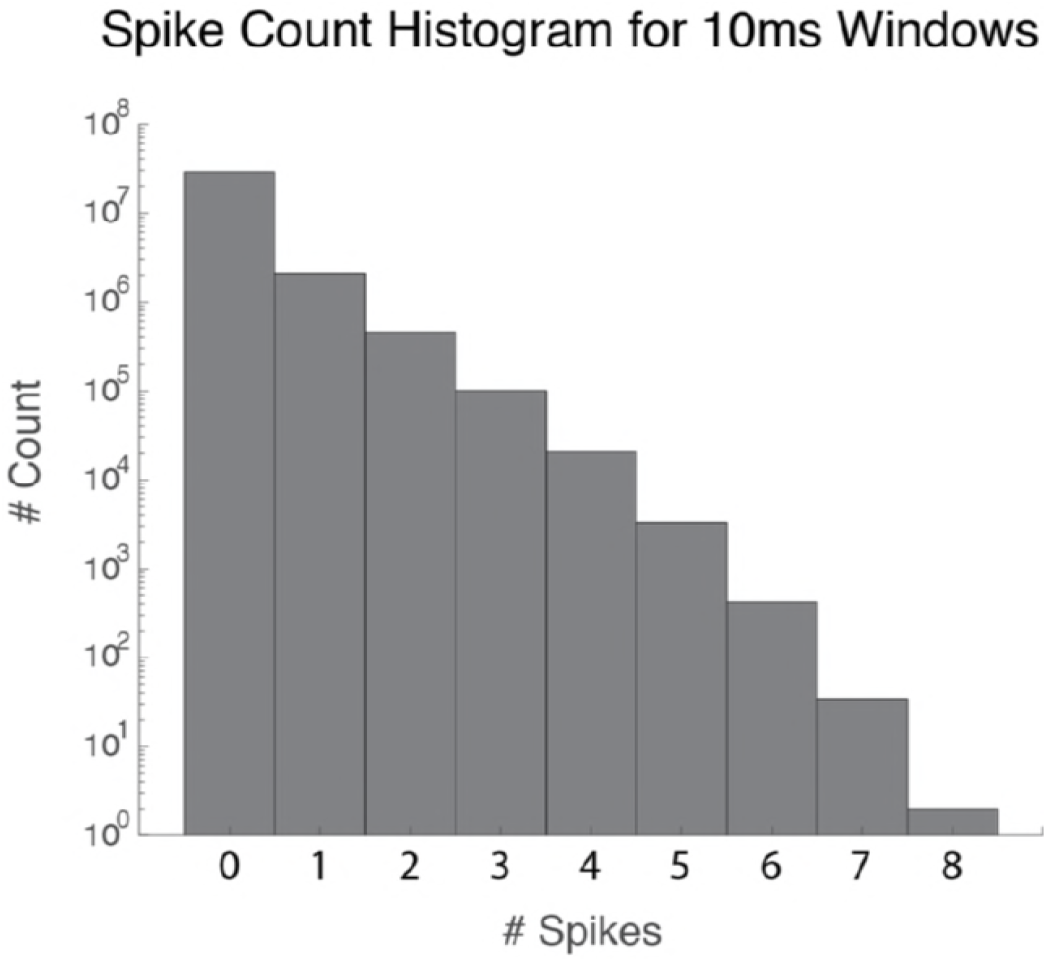
Distribution of Spike Counts in 10 ms Bins. This distribution is shown as the number of time bins across all 404 neurons, all stimuli and all time points that had 0, 1,… 8 spikes. Not a single neuron had more than 8 spikes in a 10 ms bins andhigh spiking events were very rare (only one 10ms bin with 8 spikes over all neurons, all time bins and all stimuli). The average number of spikes per 10 ms bin was 0.108.

**Supplemental Figure 3.**
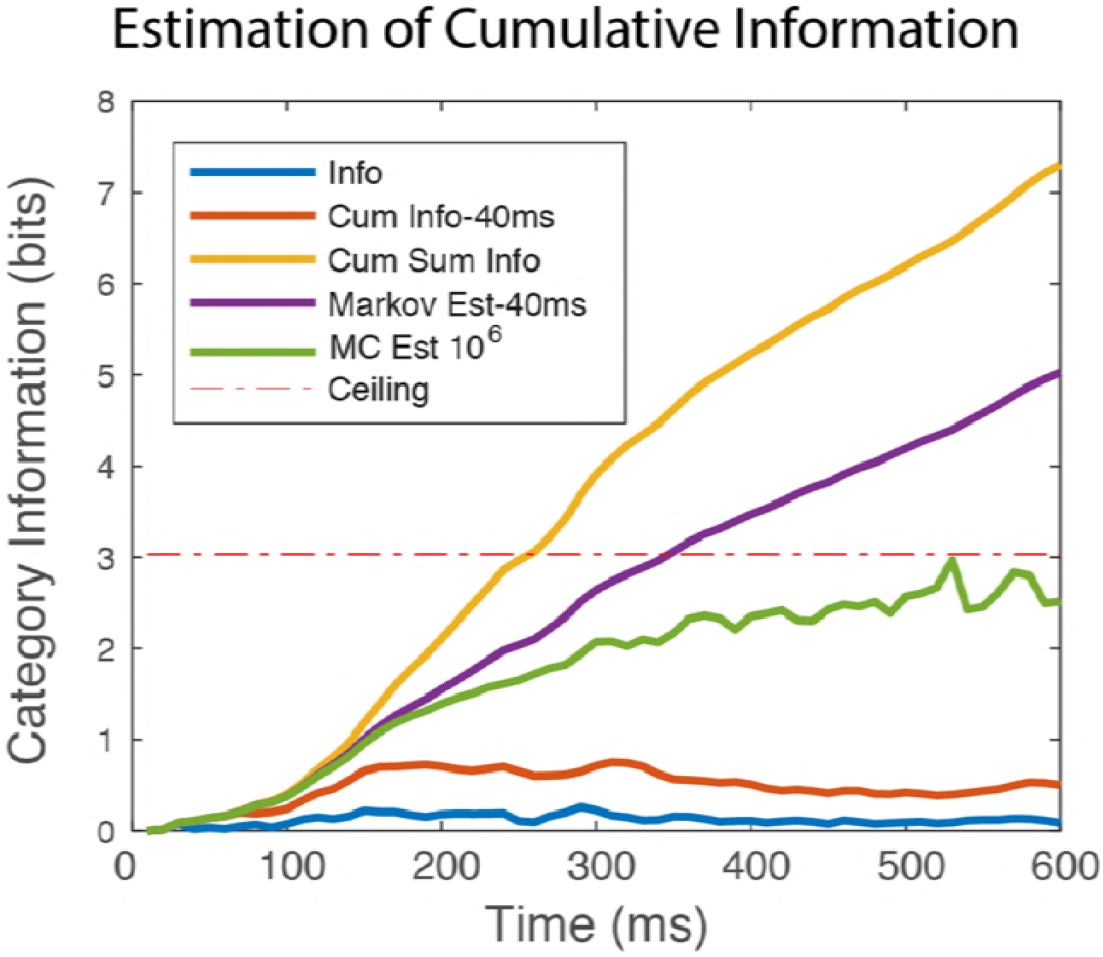
Estimation of the Cumulative Information. Three methods were tested for the estimation of the Cumulative Information (see Methods): a Markov chain approximation of variable order up to 4 or 40 ms (purple line), the exact information in 4 bins (40 ms) evaluated in running windows (red line), and a Monte Carlo estimation (green line). For comparison, the instantaneous time-varying information in 10 ms windows (blue line) and the integral of that information (yellow line) are also shown. The ceiling value corresponds here to the log2(N_cat_=9) because this example is showing the categorical cumulative information of the neuron. The Markov chain overestimates the cumulative information while the running window of 40 ms underestimates the cumulative information. The information values plotted here were obtained from the neural data of Example Neuron 1 shown in Fig. 3 and also in Sup. Fig. 4. This high-firing, high-information neuron allowed us to verify that the calculations were correct around ceiling values.

**Supplemental Figure 4.**
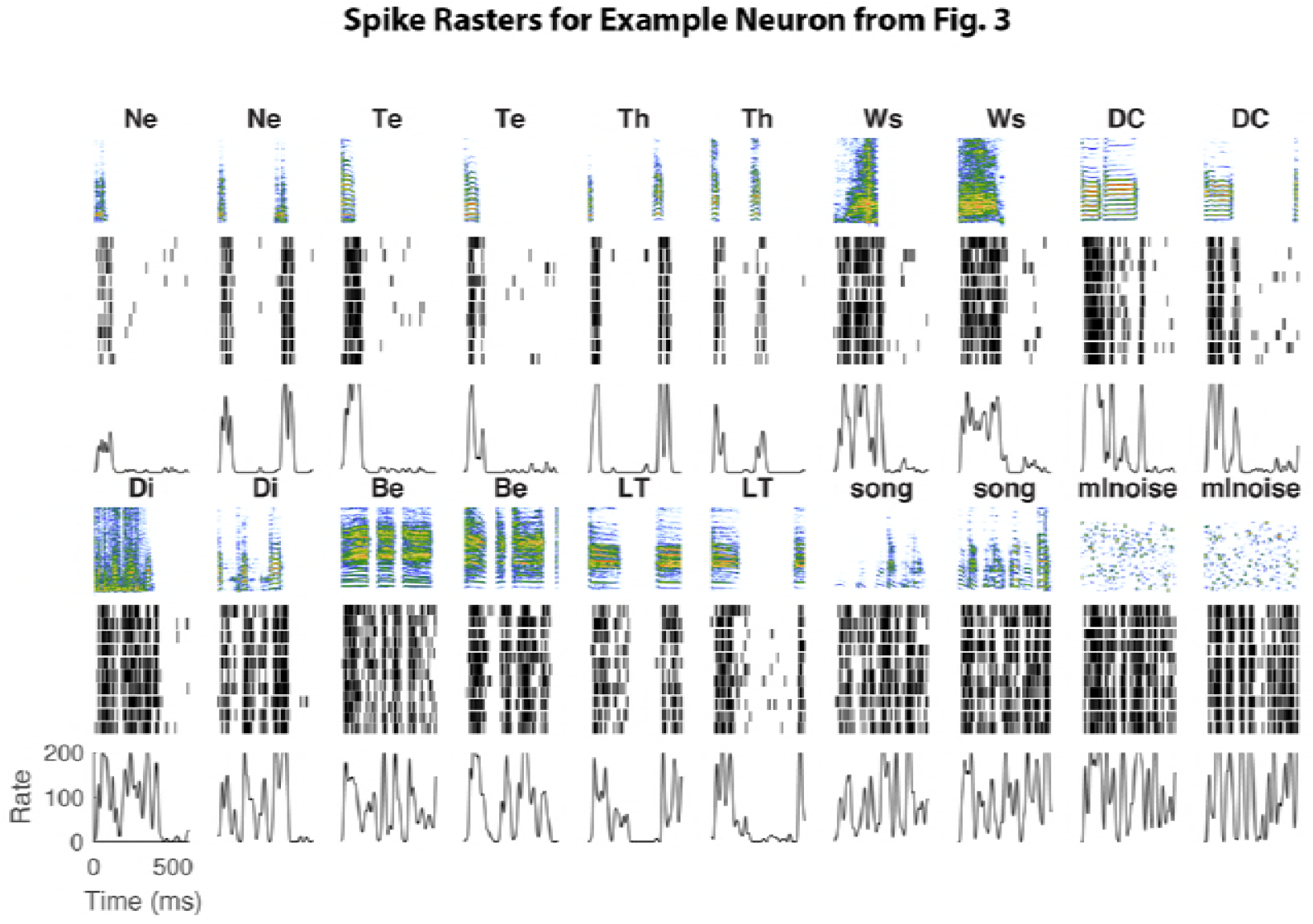
Example Spike Rasters for Example Neuron 1. The spectrogram of two (randomly chosen) stimuli from each stimulus category are shown with the corresponding spike rasters for 10 trials and a smoothed PSTH for the example neuron shown in Fig. 3. This neuron had a very high stimulus driven firing rate and responded to all sound stimuli. The mlnoise stimulus is modulation limited noise: white noise that is low-pass filtered in amplitude and spectral modulations. This stimulus was used here to search for auditory regions but the responses to these synthetic sounds were not included in these analyses.

**Supplemental Figure 5.**
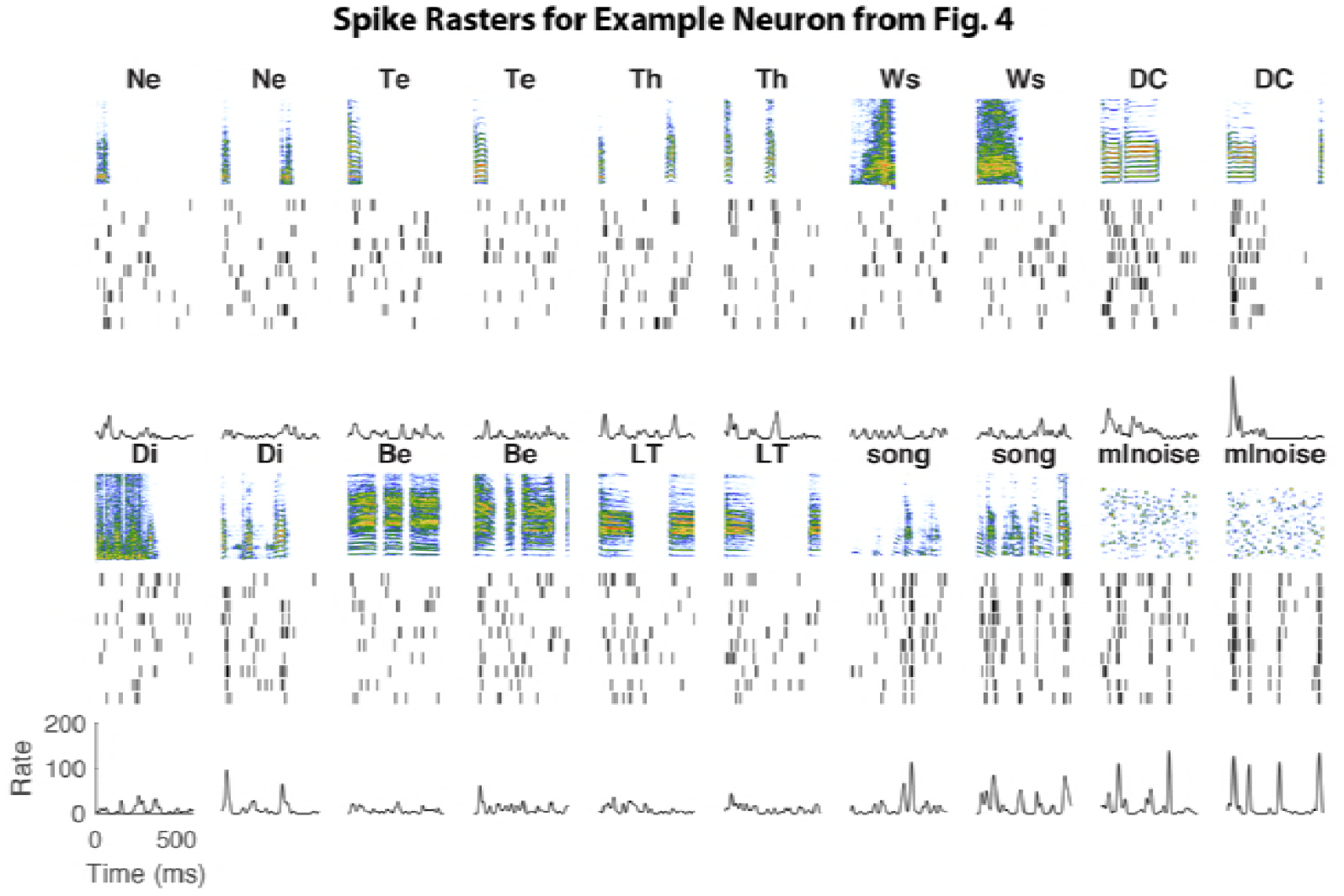
Example Spike Rasters for Example Neuron 2. As Sup. Fig. 4 but for the Example Neuron 2. Example Neuron 2 also shown in Fig. 4 of the main paper is selective for Distance Calls (DC).

**Supplemental Figure 6.**
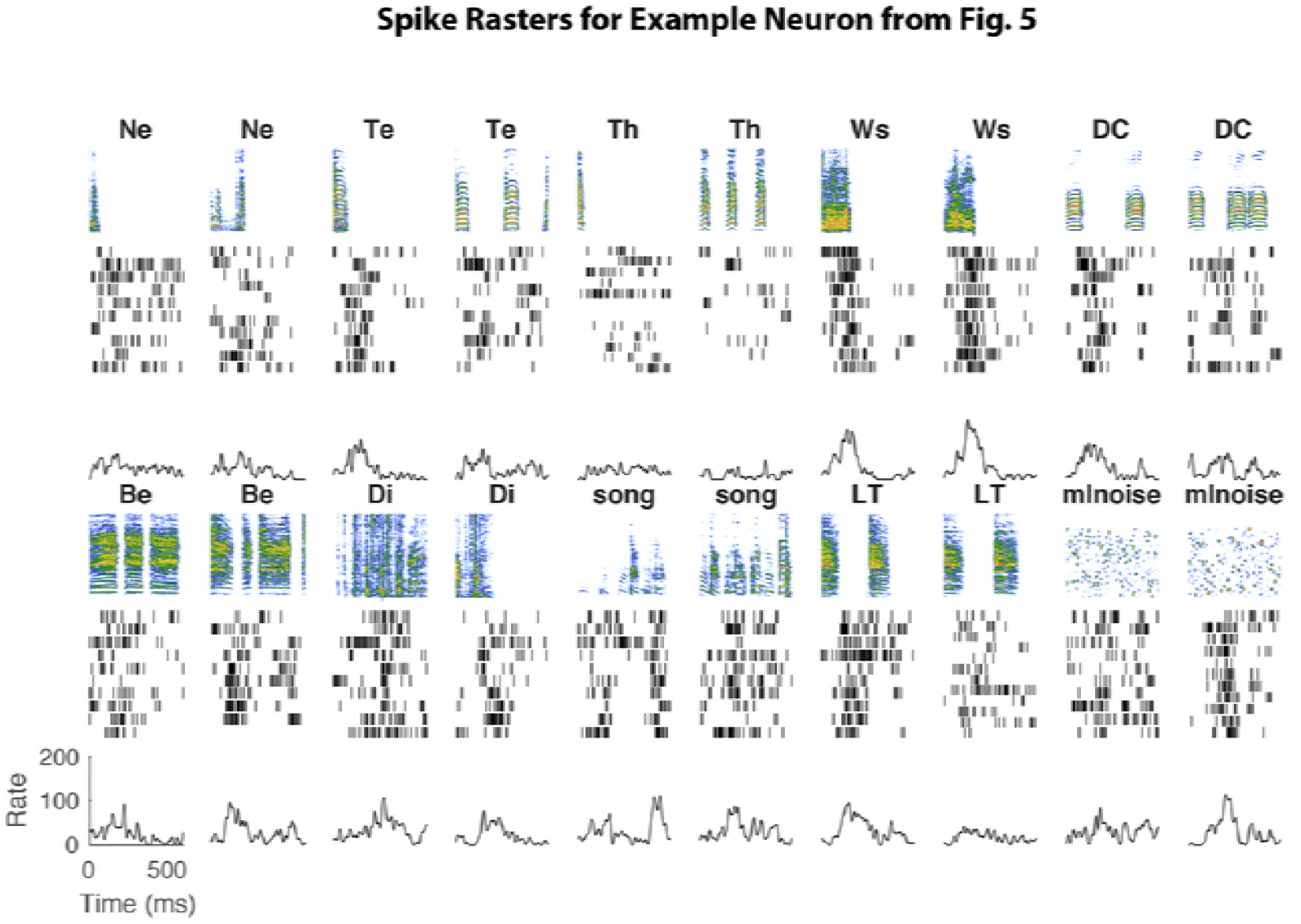
Example Spike Rasters for Example Neuron 3. As Sup. Fig. 4 but for the Example Neuron 3. Example Neuron 3 also shown in Fig. 5 of the main paper is selective for aggressive calls or Wsst Calls (Ws).

